# Behavioral deficits and exacerbated hemodynamics during lifespan of a mouse model of late onset Alzheimer’s disease expressing humanized APOEε4 and Trem2*R47H

**DOI:** 10.1101/2025.04.20.649712

**Authors:** Misha Izydorczak, Maggie Oumov, Mansiben V. Udhwani, Faysal Fostok, Guillermo Coronas-Samano, Basavaraju G. Sanganahalli, Peter Herman, Fahmeed Hyder, Justus V. Verhagen

## Abstract

Alzheimer’s Disease (AD) poses a significant global health challenge, being the most prominent cause of dementia with prevalence increasing as the population ages. While the majority of AD cases are late-onset (LOAD), current animal models predominantly represent the more aggressive, faster progressing early-onset AD (EOAD), limiting their ability in assessing early biomarkers and gaining deeper understanding of LOAD progression. This study explores a promising translatable model, the APOE4.TREM2 mouse, which combines the APOE4 allele and the Trem2 p.R47H mutation, both linked to increased AD risk in the human population. We performed behavioral phenotyping and measured hemodynamics in dorsal olfactory bulbs (dOB) during odor stimulation of the APOE4.TREM2 mouse line. Experimental evidence of olfactory dysfunction prior to clinical symptoms suggests the opportunity of utilizing smell testing and fMRI as tools for screening of AD, both for preclinical and clinical studies. Here we assess and confirm the translatability of the APOE4.TREM2 mouse LOAD model, reporting exacerbated anxiety, deficits in odor-based foraging and spatial memory, and exacerbated odor-evoked dOB intrinsic responses in an age-dependent manner.

## INTRODUCTION

Alzheimer’s Disease (AD) is an incurable neurodegenerative condition affecting millions of people worldwide and is known to be a leading cause of dementia in the aging population. Medical advancement over the years has substantially prolonged the human life span, however this has encouraged age-dependent diseases such as late onset AD, with prevalence rates increasing to 10% after the age of 65 and surpassing 40% by the age of 85 [1]. AD is classified as either familial (fAD) or sporadic (sAD) and as early onset (EOAD) or late onset (LOAD). 90% of patients with AD present with etiologically undetermined LOAD [2], yet of the approximate 168 animal models in existence the majority are known to emulate EOAD, overexpressing human genes related to fAD involved in the production of amyloid plaques and neurofibrillary tangles [3]. In contrast, genetic predisposition to LOAD has greater complexity, involving numerous genes linked to elevated risk of varying degrees [4]. It is critical to further investigate animal models that focus on the risk factors of LOAD and show clear penetrance to find early translatable biomarkers of AD. More than 20 risk factors for LOAD have been identified through genetic investigations, highlighting the pleiotropic nature of sAD [5].

The strongest genetic risk factors to developing LOAD are linked to polymorphisms of triggering receptor expressed on myeloid cells 2 (TREM2) and the apolipoprotein E 4 (APOE4) [6]. NIH recognizes the need for better animal models indicated by the large Model Organism Development and Evaluation for Late-onset Alzheimer’s Disease (MODEL-AD) effort to generate and screen new mice [7]. A promising model in a B6 mouse line combined the APOE4 and the Trem2 p.R47H mutation, where each mutation confers about a threefold risk for AD in humans that are heterozygous for either allele. This model and its wild type control are available from Jackson Laboratory (#028709 and #000664, respectively). It is critical that this APOE4-Trem2 line is scrutinized phenotypically, as it is the foundational background for introducing additional sAD associated genes [7].

APOE is a primary lipoprotein highly expressed in the brain that plays a significant role in the transport of cholesterol in the bloodstream affecting normal brain function and is associated with neurodegenerative diseases such as LOAD. The APOE gene exists in three major isoforms (ε2, ε3, ε4), and the ε4 variant has consistently been associated with an increased risk of AD. Unlike APOE2 and APOE3, APOE4 contributes to AD pathogenesis through various mechanisms that are not fully understood, but include amyloid beta (Aβ) aggregation, tau pathology, synaptic dysfunction, and brain atrophy. The APOE4 allele is the strongest genetic risk factor for Alzheimer’s disease [8].

Although APOE4 differs in only one amino acid (Arginine instead of Cystine) from APOE3, it has been recognized as an important genetic risk variable in neurological disorders, including LOAD [9]. APOE3 has a neutralizing effect such as preventing negative downstream molecular signaling and changes such as tau protein phosphorylation, promotes synaptic plasticity and Aβ clearance, and protects the brain against neurodegeneration [10]. By contrast, APOE4 inhibits neurite outgrowth, induces tau phosphorylation, disrupts neuronal cytoskeleton, and increases Aβ deposition [11].

TREM2 belongs to a family of transmembrane proteins encoded by genes on chromosome 6 [12]. These immune receptors are expressed on the surfaces of microglia, macrophages, and dendritic cells [13]. TREM2 is expressed exclusively by myeloid cells in the brain, and has been a target for extensive research in AD pathogenesis [14]. Upon injury they proliferate and migrate to damaged sites, mounting an immune response. TREM2 regulates inflammatory responses by coupling with DAP12, expressed by myeloid (and NK) cells, to initiate signaling, and either amplifies or attenuates toll-like receptor (TLR) induced signals [12]. Ongoing AD genetic studies have identified multiple AD-associated polymorphisms, with the R47H variant in a TREM2 missense mutation being associated with the greatest risk. The R47H variant demonstrated a deficiency in ligand binding and showed the greatest defect in TREM2 activation/expression ratio [15].

Neuroimaging studies in humans have shown that even cognitively normal APOE4 carriers develop vascular, metabolic, and structural deficits decades before the aggregation of Aβ and neurofibrillary tau tangles [16]. Interventions that can restore these deficits to normal could be critical to prevent the development of AD-related neuropathology and cognitive impairment [17]. However, before interventions in humans can be done, more information about the development and molecular mechanisms underlying these deficits is needed. Animal models, especially transgenic mouse models, that develop LOAD-like pathology and impairments, could provide flexible, rigorous, and systematic approaches for understanding LOAD development.

Clinical symptoms of AD involve progressive mental, behavioral, and functional impairments. Pathophysiological changes vary depending on the disease progression and on the continuum of disease severity. The entorhinal cortex and hippocampus are two brain regions intricately linked to memory and closely associated with the onset of AD-related cognitive impairments. Early signs of AD often include deficits in episodic and spatial memory, highlighting their significance as initial cognitive indicators of AD [18]. Additionally, there are numerous noncognitive behavioral symptoms associated with AD such as anxiety, depression, and sleep disturbances [19, 20]. Depression symptoms may appear before the onset of AD [21] and people who have previously been diagnosed with depression are more likely to experience a reoccurrence over the course of AD [22].

Similar to behavioral indicators, olfactory dysfunction is another early marker of dementia and has been used as an early indicator of neurodegenerative disorders such as AD with prevalence as high as 100% [23]. Early impairment of central olfactory regions like the entorhinal cortex and hippocampus link the olfactory system to AD’s initial manifestation, serving as an early indicator of AD related dysfunction [23]. Olfactory deficits are known to be associated with a four to fivefold increased risk of developing AD for patients already presenting with mild cognitive impairment (MCI) [24]. Therefore, odor-associated tasks and optical intrinsic imaging of the olfactory bulb during odor-evoked activation have great promise toward increasing our understanding of LOAD development and for identifying translatable early biomarkers.

Our study characterizes the age-dependent changes associated with APOE4.Trem2 in the context of LOAD, employing a dual approach of behavioral and hemodynamic analysis to build on prior multipronged assessments by the MODEL-AD consortium. Mice expressing the humanized APOE4 allele showed decreased plasma lipoprotein levels and age-dependent decrease in glucose metabolism [7]. LOAD’s progression is substantially less aggressive and has dependence on many more genetic risk factors, as well as on aging, inflammation and life style, compared to EOAD. Previous findings indicate the APOE4.TREM2 model does not produce severe, strong LOAD phenotypes, even late into life, important for a better understanding of aging in relation to AD onset and progression [7]. However, it has been reported [25] that introducing the R47H mutation, as used in this study, results in the generation of a distinctive murine splice site, causing a roughly 50% reduction in TREM2 protein and a 20% decrease in Trem2 transcription. Deciphering the effects on microglial response due to the mutation and decreased expression is challenging without accurate representation of amyloid plaques. Regardless, APOE4 has shown to consistently replicate the biology observed in other APOE mouse models, and, importantly, in humans [26].

This study assesses the translatability of the APOE4.TREM2 model through behavioral phenotyping, in addition to alterations in hemodynamic response, in order to establish early biomarkers for LOAD. Our evaluation of AD-related behaviors encompasses seven tests: the Morris Water Maze for spatial memory, the habituation/ dishabituation task for olfactory memory and function, the hidden cookie test for odor navigation and foraging, the open field test for anxiety, the sucrose preference test as an anhedonia/depression assessment, the nestlet building test, and the sleep/wake cycle. Odor-evoked intrinsic optical imaging of the dorsal olfactory bulb (dOB) in freely breathing mice allowed for visualization and quantification of vascular activity throughout varying stages of disease. Previous fMRI performed on patients with MCIs revealed compensatory hemodynamic responses while performing cognitive tasks. Vascular hyperactivity was found compared to healthy aged adults, specifically in the right pre-frontal cortex, and was especially evident in patients performing the Stroop test [24]. Our study suggests that fMRI odor-response imaging may be an effective diagnostic method for early LOAD detection.

## METHODS – BEHAVIORAL ASSAYS

### Subjects

In this study we compared the behavior of aged B6(SJL)- *Apoe^tm1.1(APOE*4)Adiuj^ Trem2^em1Adiuj^*/J (n=23), Strain #028709, with WT mice (n=27), Strain #000664 from the Jackson Laboratory (www.jax.org). The mice were divided into three age groups and run in 3 batches (**Table 1**). **Table 1** lists the number of APOE4.TREM2 and WT mice used in these behavioral studies by genotype, gender, and age. **Table 2** shows the number of mice by sex, age and genotype for each behavioral test. Mice had a rest period of at least 24 hours between the tests. A 12-hour/12-hour inverted light cycle with lights out at 9:00 am was used in the vivarium. The mice were fed ad libitum chow (Harlan 18% protein rodent diet) and were housed separately in polycarbonate cages (12 x 12 x 25 cm) with controlled humidity (40%) and temperature (22°C). Guidelines established by the National Institute of Health were followed for the care of all animals (2011). The experimental protocols were approved by the Institutional Animal Care and the John B. Pierce Laboratory (JV1-21). The John B. Pierce Laboratory is AAALAC accredited.

**Table 1.**
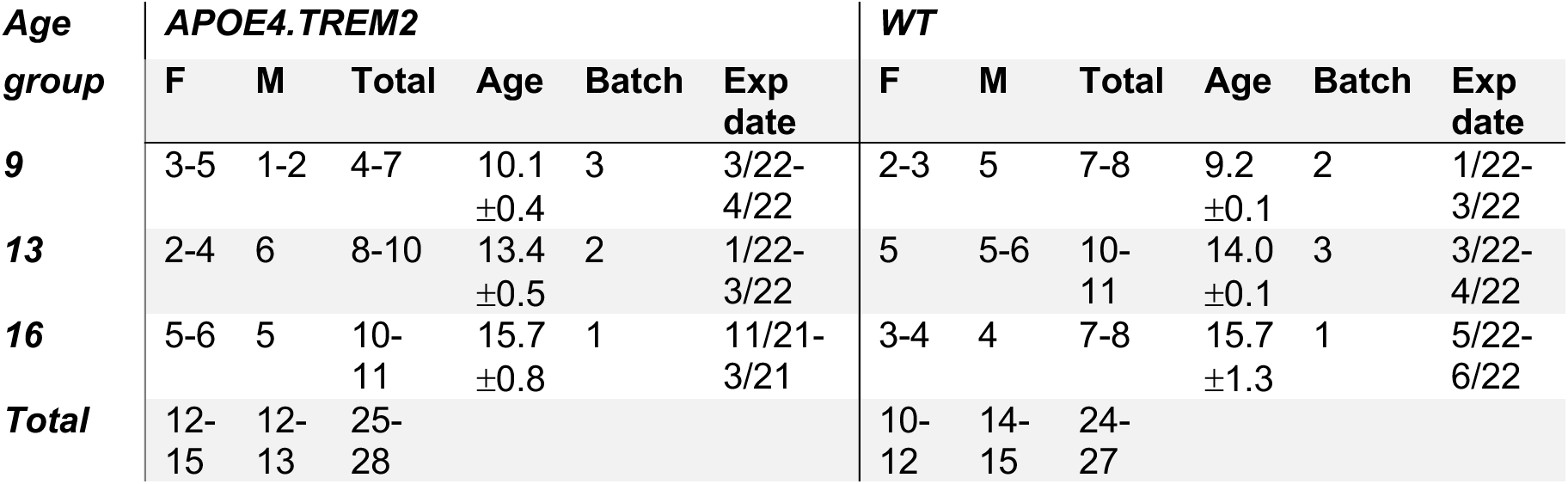
The number of mice in each genotype, gender, and age used for the behavioral study. Their experimental batch number and date range of experiments are also indicated.

**Table 2.** The number of mice in each genotype, gender, and age of the APOE4.TREM2 and WT control mice used for the behavioral study. Boldfaced numbers have missing data due to incomplete trials. SW: sleep/wake cycle activity; SPT: sucrose preference test; NB: Nest building; MWM: Morris water maze; OFT: open field test; HC: hidden cookie test; HaxHa: odor habituation/dis-habituation test.

| Age group | Batch | Genotype | SW |  | SPT |  | NB |  | MWM |  | OFT |  | HC |  | HaxHa |  |
| --- | --- | --- | --- | --- | --- | --- | --- | --- | --- | --- | --- | --- | --- | --- | --- | --- |
|  |  |  | F | M | F | M | F | M | F | M | F | M | F | M | F | M |
| 9 | 2 | WT | 3 | 5 | 3 | 5 | 3 | 5 | 2 | 5 | 3 | 5 | 3 | 5 | 3 | 5 |
| 9 | 3 | APOE4.TREM2 | 5 | 2 | 5 | 2 | 3 | 1 | 4 | 2 | 5 | 2 | 4 | 2 | 5 | 1 |
| 13 | 3 | WT | 5 | 6 | 5 | 6 | 5 | 6 | 5 | 5 | 5 | 6 | 5 | 6 | 5 | 5 |
| 13 | 2 | APOE4.TREM2 | 4 | 6 | 4 | 6 | 4 | 6 | 2 | 6 | 4 | 6 | 4 | 6 | 4 | 6 |
| 16 | 1 | WT | 4 | 4 | 4 | 4 | 4 | 4 | 3 | 4 | 4 | 4 | 4 | 4 | 4 | 4 |
| 16 | 1 | APOE4.TREM2 | 5 | 5 | 5 | 5 | 6 | 5 | 6 | 5 | 6 | 5 | 6 | 5 | 6 | 5 |
| All 9-16 | 1-3 | WT | 12 | 15 | 12 | 15 | 12 | 15 | 10 | 14 | 12 | 15 | 12 | 15 | 12 | 14 |
| All 9-16 | 1-3 | APOE4.TREM2 | 14 | 13 | 14 | 13 | 13 | 12 | 12 | 13 | 15 | 13 | 14 | 13 | 15 | 12 |

### Statistical analysis

Statistical analyses were performed in Prism version 10.4.2 for Mac OS (GraphPad Software, Boston, Massachusetts USA, www.graphpad.com). Means are reported as means±SEM. Post-hoc tests were Bonferroni corrected for multiple comparisons.

### Behavioral tracking with Noldus Ethovision

All behavioral tests tracking movement (Habituation/cross-habituation, Open Field Test, Hidden cookie test, Morris Water Maze) were assessed by Noldus Ethovision (EthoVision XT, version 10.1, Noldus Information Technology B.V., Wageningen, The Netherlands). A video camera (Basler Gen1cam, Resolution 1280*1024) connected to a Noldus computer system was placed above the test arena.

### Habituation/cross-habituation test (HaXha)

This olfactory-dependent behavioral task was performed in a sealed semi-transparent white acrylic box (26x38x16cm) in which a cotton-tipped wood applicator (8cm long, Dynarex MS-50330) was placed in the center of the box and presented to the mouse, while retained by a 54x54x4mm applicator holder. We added one of 3 odorants to the applicator: mineral oil (MO, control), amyl acetate (AA) (diluted 1% in MO), phenyl ethanol (PE) (1% in MO), and social odor (S, SO) (obtained by swabbing the cage of non-experimental CRE-OMP female mice). A total of 12 trials were done per mouse, where each odorant was presented three times in succession per session to yield the following order: MO1-3, AA1-3, PE1-3 and SO1-3. Each trial consisted of a 2 min odorant exposure and an interval of 1 min between stimuli. The time spent smelling the cotton tip was registered using a camera mounted at the ceiling of the box, and Ethovision identified the orientation and scored the behavior of the animal. Smelling was defined as being oriented toward the applicator tip while the nose is within 2 cm of it, which was validated by us in [27]. This test evaluates if mice can spontaneously recognize a novel odorant stimulus by spending more time smelling the applicator (cross-habituation phase, trial 3 vs new odor 1), as opposed to the time that the mice spent on each stimulus (habituation phase, trial 1 vs 3 of same odor).

### Sucrose preference test (SPT)

SPT was divided into Day 1 and 2 measuring the volume of water and sucrose solution consumption. Our sucrose preference test was based on [28]. The mice were placed in regular home cages (22.5 x 16.7 x 14 cm) and had access to two drinking tubes, one filled with 20 ml tap water, and the other with 20 ml 10% sucrose (Sucrose, Lot 18005800, American bio) diluted in deionized water. The location of the tubes was randomized for two consecutive days. Six hours the free consumption of water and 10% sucrose solution took place in the presence of *ad libitum* food during the dark phase. Sucrose preference was the percentage of the total amount of liquid drunk in 6 hours that was sucrose.

### Open Field Test (OFT)

The Open Field was based on [29–31] to assess spontaneous exploratory activity and anxiety-related behaviors. Anxiety and exploratory activities were evaluated by allowing mice to freely explore an open field arena for 15 min. The testing apparatus was a classic open field (i.e., a white PVC square arena, 50 x 50 cm, with walls 40 cm high). An Ethovision-connected video camera was placed above the box. Each mouse was placed individually on the center of the arena and the behavior was monitored by the video tracking system (Noldus Ethovision). The central area was arbitrarily defined as a square of 35 x 35 cm (half the total area). The ethological measures included the frequency and duration spent at each area (center and periphery) and locomotor activity.

### Hidden cookie test (HCT)

This test assesses deficits in olfactory guided foraging skills by measuring the latency of the mice to find a buried cookie in one of the corners of their cage. Mice were familiarized with the chocolate cookie (Chips Ahoy!) for a week before the test. A quarter of chocolate cookie (∼2.5 g) was placed every 48 hours for three times in the home cage on the top of the bedding. Mice received food and water ad libitum. After a week of the familiarization phase, mice were deprived of the cookie for 3 days before the test. On the first day of the test, mice were food restricted for 6 hours and individually caged with 3 cm of clean bedding. Initially, a piece of cookie was buried (2 cm deep) in one corner of the cage and the latency to finding the cookie was recorded using Noldus Ethovision. An hour and a half later, another piece of cookie was buried in the same corner as the first trial, and the behavior was tracked and analyzed. At the end of the second trial, the mice were taken to the animal room. The next day, mice were placed in the experimental cages and the piece of cookie was buried in a corner different from the last day, and this was tested twice following the same protocol as the first experimental day.

### Morris Water Maze test (MWMT)

Spatial reference learning and memory were evaluated in a water maze adapted from that previously described by Morris and collaborators [32] and was based on [29–31, 33]. The test was performed in a circular stainless steel pool of 90 cm in diameter and 40 cm height, filled with 20 cm of water tainted with a non-toxic white paint (acrylic paint, 20503 white, Apple Barrel). As it is known that the standard Morris water maze can interfere with physiology, likely due to the stress of heat loss, we opted to refine this method to be less stressful by not maintaining it at room temperature (21±2°C), but instead between room and core body temperature (37°C), i.e. at 29±2°C. The pool was virtually divided into four equal quadrants, labelled north–south–east–west. The Ethovision camera was mounted above the maze.

During training, a platform (10 cm in diameter and made of transparent acrylic plastic) was submerged 1 cm below water surface and was placed at a fixed location in one of the quadrants. If the mice were not able to reach the platform within 120 s, they were guided to the platform where they had to stay for 30 sec before being returned to their home cage for 30 sec. All mice were given four trials per day, once from each quadrant, for four consecutive days. The start position was randomized among four quadrants of the pool. During training trials, the latency to reach the escape platform and the path length were measured.

A probe trial was performed 24 h after the last day of training. During the probe trial, mice were allowed to swim in the pool without the escape platform for 120 s. The latency to reach the platform (s), swim distance (cm), and swim speed (cm/s) were recorded using Ethovision. During the probe trial, performance was expressed as the percentage of time spent in each quadrant of the MWM, and the number of crossings through the position where the platform used to be during acquisition (using 15 cm diameter target area).

### Nestlet building task (NBT)

After at least a 24 h rest period, mice were housed individually and tested for nest building (adapted from [34, 35]). Two hours prior to the onset of the dark phase of the light cycle, individual cages were supplied a nestlet made of pressed cotton square (Ancare, UK agent, Lillico). The next morning (∼24 h later) cages were inspected for nest construction. Pictures were taken prior to evaluation for documentation. Nestlet nest construction was scored by 3 investigators blind to genotype using the system of Deacon (please see [34] for detailed scoring standard). Briefly, in this 5-point scale, 1 indicates a >90% intact nestlet, whereas a 5 indicates a nestlet torn >90% and a clear nest crater.

### Sleep/awake task

Piezo foil (PVDF)-based circadian rhythm scoring of sleep/wake rhythm was used as established in [36] and used in several AD studies [37–39]. Mice were individually housed for 48 hours in clean plastic cages (Signal Solutions LLC, www.sigsoln.com, Lexington, KY) with approximately 5 mm of corn cob bedding lining the floor on top of the PVDF foil and left with an inverted light cycle (as in the colony) in a separate dedicated room. Activity was continuously recorded without disturbance for the 48 hours and was analyzed for the circadian rhythm of wakefulness and sleep using the Signal Solutions software.

## METHODS – OLFACTORY BULB OPTICAL IMAGING

### Subjects

This portion of the study uses the same strains as the behavioral study and were housed under the same conditions. Here we evaluate the difference in neurovascular coupling between aging B6(SJL)-Apoe^tm1.1(Apoe*4)Adiuj^ Trem2^em1Adiuj^/J (n= 17) versus WT (n= 21). **Table 3** shows the number of mice used for this study by age, gender, and genotype. Guidelines established by the National Institute of Health were followed for the care of all the animals (2011). The experimental protocols were approved by the Institutional Animal Care and the John B Pierce Laboratory. The John B Pierce laboratory is AAALAC accredited.

**Table 3.**
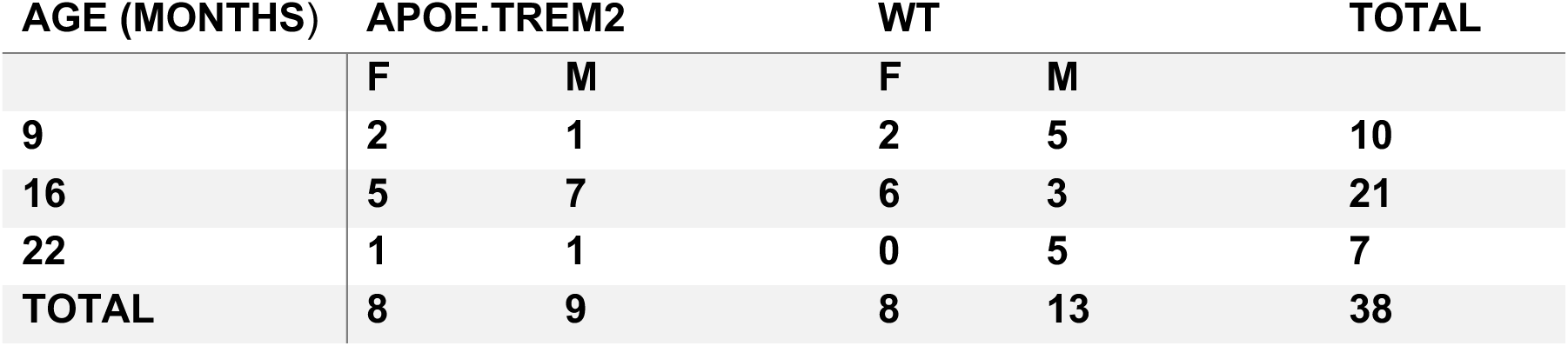
The number of mice in each genotype, gender, and age range of the APOE4.TREM2 and WT control mice used for the imaging study.

### Headplate Fixation and Olfactory Bulb Exposure

Mice were anesthetized with isoflurane (4% for induction, 1.5-3% for maintenance) followed by injections of Atropine (0.04mg/kg, IP) and Ethiqa XR (3.25mg/kg, SC). Body temperature was maintained at 37°C. An incision was made down the midline of the skull between the eyes exposing the dorsal surface of the bulbs, followed by the administration of topical bupivacaine. Custom headplates were placed and secured with MetaBond.

Once the animal was placed in the stereotax a dental drill was used to thin the bone and expose the dorsal side of the olfactory bulbs (dOB). Sterile saline was repeatedly used to saturate the skull to assess the optical clarity. The skull over the bulbs, as well as additional exposed skull and skin edges, were sealed with Krazy Glue for transparency. Next, Carprofen (5mg/kg, SC) was injected. Mice are kept isolated in recovery cages while body temperature was supported until awake and then given 4-8 days to recover prior to imaging.

### Intrinsic Imaging

Mice were anesthetized with isoflurane (4%) followed by injections of Dexdomitor (0.5mg/kg, IP), Ketamine (75mg/kg, IP), and Atropine (0.04mg/kg, IP) with body temperature maintained at 37°C. Mineral oil was put on the bulbar window to enhance optical clarity. Intrinsic optical signals from the dOB were recorded using a CCD camera (Redshirt Imaging LLC, Decatur, GA, USA) with 256×256 pixel resolution, at a frame rate of 7 Hz. The epifluorescence macroscope used was a custom-made tandem-lens type [40]. Imaging lenses were prime Nikon F-mount (ccd lens: 135mm f/3.5, used at f/8; object lens: 135mm f/3.5, used at f/3.5 or 80mm at f/5.6). A high-power LED with 590nm spectral peak (M590L4, Thorlabs, Newark, NJ), chosen for its isosbestic wavelength of oxy- and deoxy-hemoglobin to specifically measure blood volume (BV), was aimed via an optical fiber at the dOB, driven by a T-Cube LED Driver (LEDD1B, Thorlabs, Newark, NJ).

The flow dilution olfactometer output was presented through Teflon tubes that converged to a mask positioned to isolate the nose of the mouse. Nine trials using ethyl butyrate (EB) at three increasing concentrations (0.3%, 1.0%, and 3.0% of saturated vapor pressure, respectively) were carried out after the initial three control trials with no odor presented. Odor was presented for 4 seconds, 8 seconds into the start of the trial. A 180 second interval was used between each trial allowing OB activity to ensure return to baseline. Once the EB trials were completed, we used a 5-minute time interval to clear out all remaining odor. The same process was repeated for additional controls and methyl valerate (MV) at the same dilutions. After all trials were completed, mice were administered Antisedan (1mg/kg, SC) and were allowed to recover while supporting body temperature.

### Regions of Interest (ROI) Selections

MATLAB (The Mathworks Inc., V 2023) was used for dOB arterial region selection, by selecting pixels based on the consistency of blood volume (BV) increase out of the six trials from each concentration. Three regions were evaluated, the first ROI selected pixels with BV increase one SD above baseline for 4/6 trials **(Figure 8a**), the second selected pixels with BV increase three SD above baseline for 3/6 trials **(Figure 8b**), and the third ROI being a rectangular selection with no thresholds or assumptions, thereby evaluating BV changes across a large section of each dOB **(Figure 8c**). For further validation of these regions, mice were imaged during an intravenous tail injection of Fluorescein isothiocyanate (FITC, 250mg/kg) using a high-power LED of 488nm (Thorlabs, Newark, NJ) aimed via optical fiber at the dOB, driven by a T-Cube LED Driver (LEDD1B, Thorlabs, Newark, NJ). Imaging was captured using the same imaging setup, at 25 frames/s.

### Image Analysis

Unless indicated, all optical imaging data analysis was done using custom-written *MATLAB* (The Mathworks Inc., V 2023).

## RESULTS

### Habituation-Cross-habituation test reveals no odor memory deficits in APOE4.TREM2 mice

The Habituation-Cross-habituation test aimed to determine how APOE4.TREM2 mice might display olfactory dysfunction in comparison to WT animals throughout their lifetime (**Table 1, 2**; see **Methods** for full details). This assessment involved evaluating their capacity to distinguish between a sequentially presented set of odors. Mineral oil (MO) functioned as the odorless diluent used as the control stimulus to measure baseline exploration and habituation. Phenyl ethanol (PE) served as a non-trigeminal rose-like odorant, allowing for the selective probing of the olfactory system. Amyl acetate (AA) was selected as a commonly used banana-like food odorant. Lastly, social odor (SO) was included, as previous research indicated that this stimulus elicited longer exploration times [41], thereby enhancing the sensitivity of the test. Post-hoc we performed seven Bonferroni-corrected comparisons between the first and third of each stimulus (habituation), and the third and following first presentation (cross-habituation, or “dis-habituation”).

At 9 mo, the WT mice showed an overall effect of exploration time across the odors (F_(1.35, 9.42)_ = 8.32, *P* = 0.02; one-way RM ANOVA, **Figure 1a**, **Table 1 and 2**). WT mice displayed habituation (•) between the repeated presentation of the same social odors; SO1 (“S1”, 28.7 ± 6.9 s) and SO3 (“S3”, 11.3 ± 6.2 s; ··*P* = 0.003). They also showed cross-habituation (◆) between the presentation of PE3 (1.8 ± 0.5) and SO1 (28.7 ± 6.9 s; ◆*P* = 0.04). Interestingly, the 9 mo APOE4.TREM2 mice did not show an overall effect on exploration time across the odors (F_(2.88, 14.4)_ = 2.67, *P* = 0.09; one-way RM ANOVA, **Figure 1a**). Therefore, habituation and cross-habituation were not significant. Planned post-hoc comparisons were also not significant. Additionally, we found statistical difference in the exploration time of SO1 between 9 mo WT (28.7 ± 6.9 s) and the APOE4.TREM2 group (16.4 ± 5.0 s; * *P* = 0.03), however there was no overall difference between the groups (F_(1, 143)_ = 0.84, *P* = 0.36, two-way ANOVA, **Figure 1a**).

**Figure 1.**
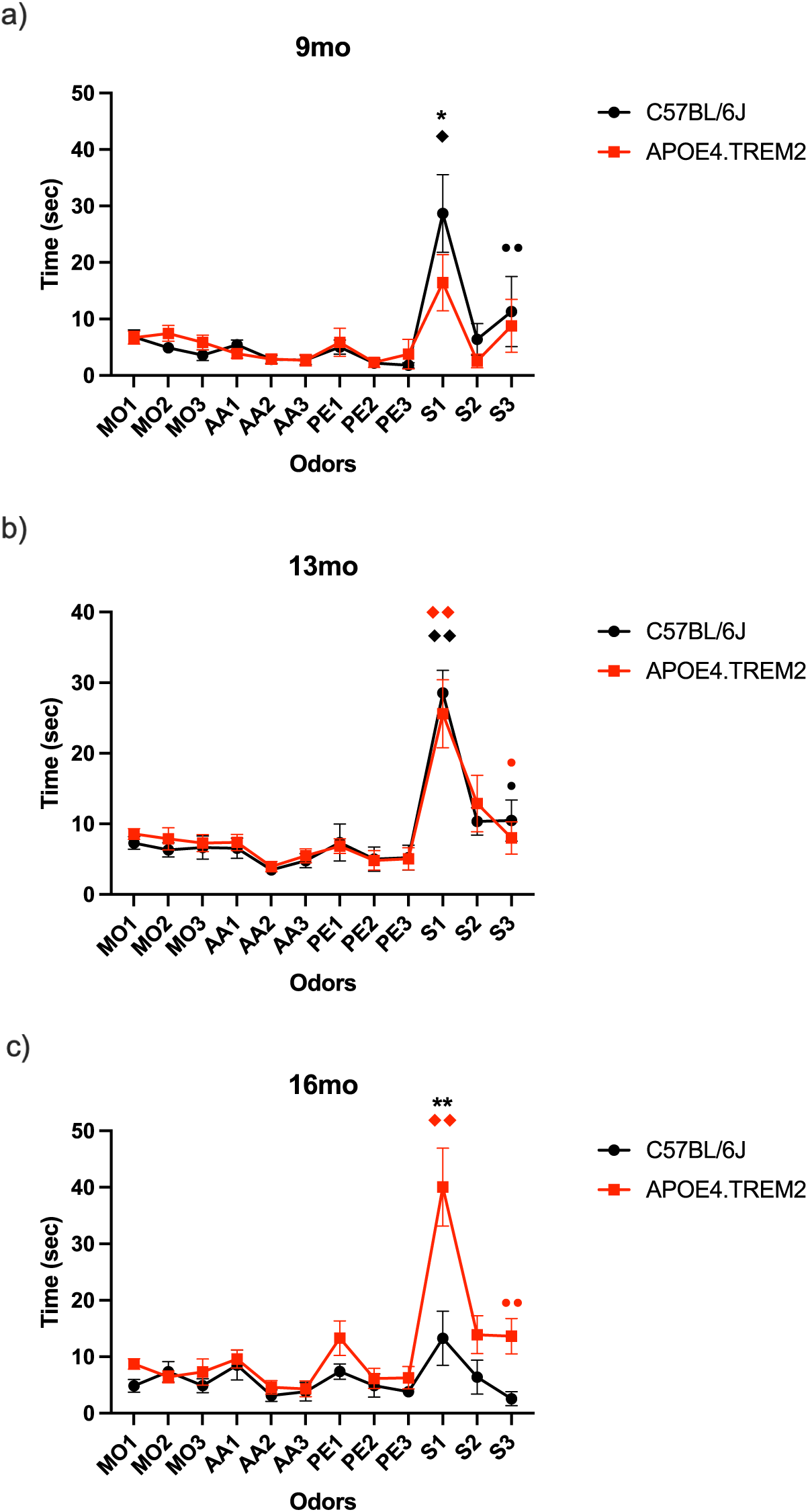
Habituation-Cross-habituation test. Data depicts mean ± SEM for the time (sec) spent by WT (black) and APOE4.TREM2 mice (red), sniffing one of the following odors: MO, PE, AA, or a social odor to test APOE4.TREM2 and WT mice’s ability to distinguish novel odors. (**A**) graphical representation of 9 mo. old WT (black) and APOE4.TREM2 (red). (**B**) Graph illustrating 13 mo. old WT (black) and APOE4.TREM2 (red) mean ± SEM odor exploration. (**C**) Graph representing 16 mo. WT (black) and APOE4.TREM2 mice (red). * p<0.05 vs WT, ** p<0.0001 vs WT. ◆p<0.05, ◆ ◆p<0.01 cross-habituation. • P<0.05, ·· P<0.01 habituation. See **Table 2** for number of mice per group.

The 13 mo, WT mice showed an effect on exploration time across the odors (F_(3.77, 33.8)_= 13.9, *P <* 0.0001; one-way ANOVA, **Figure 1b**). This group showed cross-habituation between PE3 (5.2±1.8) and SO1 (28.6±3.2, ◆ ◆P < 0.001) and habituation between SO1 and SO3 (10.5±2.9, •P = 0.02). Meanwhile, the APOE4.TREM2 group showed a stimulus effect on exploration duration (F_2.52, 22.67_ = 7.7, *P* = 0.002, one-way ANOVA, **Figure 1b**) with cross-habituation between PE3 (5.1 ± 1.6 s) and SO1 (25.6 ± 4.8 s; ◆ ◆*P*= 0.007), and habitation between SO1 and SO3 (8.0 ± 2.3 s, • *P* = 0.04). There were no overall differences between the 13 mo groups (F_(1, 216)_ = 0.03, P = 0.86), nor planned post-hoc differences.

At 16 mo, WT mice showed no effect of stimulus on exploration time (F_(1.99, 13.9)_ = 2.2, *P* = 0.15; one-way ANOVA, **Figure 1c**). No habituation or cross-habituation was observed. In contrast, the 16 mo APOE4.TREM2 mice showed an effect on exploration time across the odors (F_(2.35, 23.50)_ = 11.90, *P* = 0.0002, one-way ANOVA, **Figure 1c**). This group showed habituation between SO1 (40.0 ± 6.9 s) and SO3 (13.7 ± 3.1 s; ··*P* = 0.004). The cross-habituation was significant between PE3 (6.3 ± 2.0 s) and SO1 (40.0 ± 6.9 s; ◆ ◆*P* = 0.009). Interestingly, there was a statistical difference for SO1 between the 16 mo WT (13.3 ± 4.8 s) and the APOE4.TREM2 (40.0 ± 6.9 s; **P < 0.0001) (F_(1, 204)_ = 22.71, *P* < 0.0001, two-way ANOVA, **Figure 1c**).

These results do not suggest age-related olfactory dysfunction in the APOE4.TREM2 mice, although they did exhibit an unexpected increased novel social odor exploration with age.

### Sucrose preference task: APOE4.TREM2 mice do not show anhedonia, but lower overall fluid intake

Consumption was measured over two days in the sucrose preference task, testing the mice’s reward sensitivity for indications of anhedonia. The mice were placed in regular home cages and had access to two drinking tubes, one filled with 20 ml tap water, and the other with 20 ml 10% sucrose diluted in deionized water. Six hours of free consumption of water and 10% sucrose solution took place in the presence of *ad libitum* food. Sucrose preference was calculated from the amount of sucrose solution consumed, expressed as a percentage of the total amount of liquid ingested in 6 hours.

At 9 mo both groups obtained high preference on the first day of the test, WT 87.5 ± 2.5% and APOE4.TREM2 85.7 ± 5.3 % consumption. Preference was maintained on the second day, WT 92.5 ± 2.5 % and APOE4.TREM2 92.9 ± 1.8 %, with no statistical differences between groups (F_(1,26)_=0.05, P=0.83; two-way ANOVA, **Figure 2a**).

**Figure 2.**
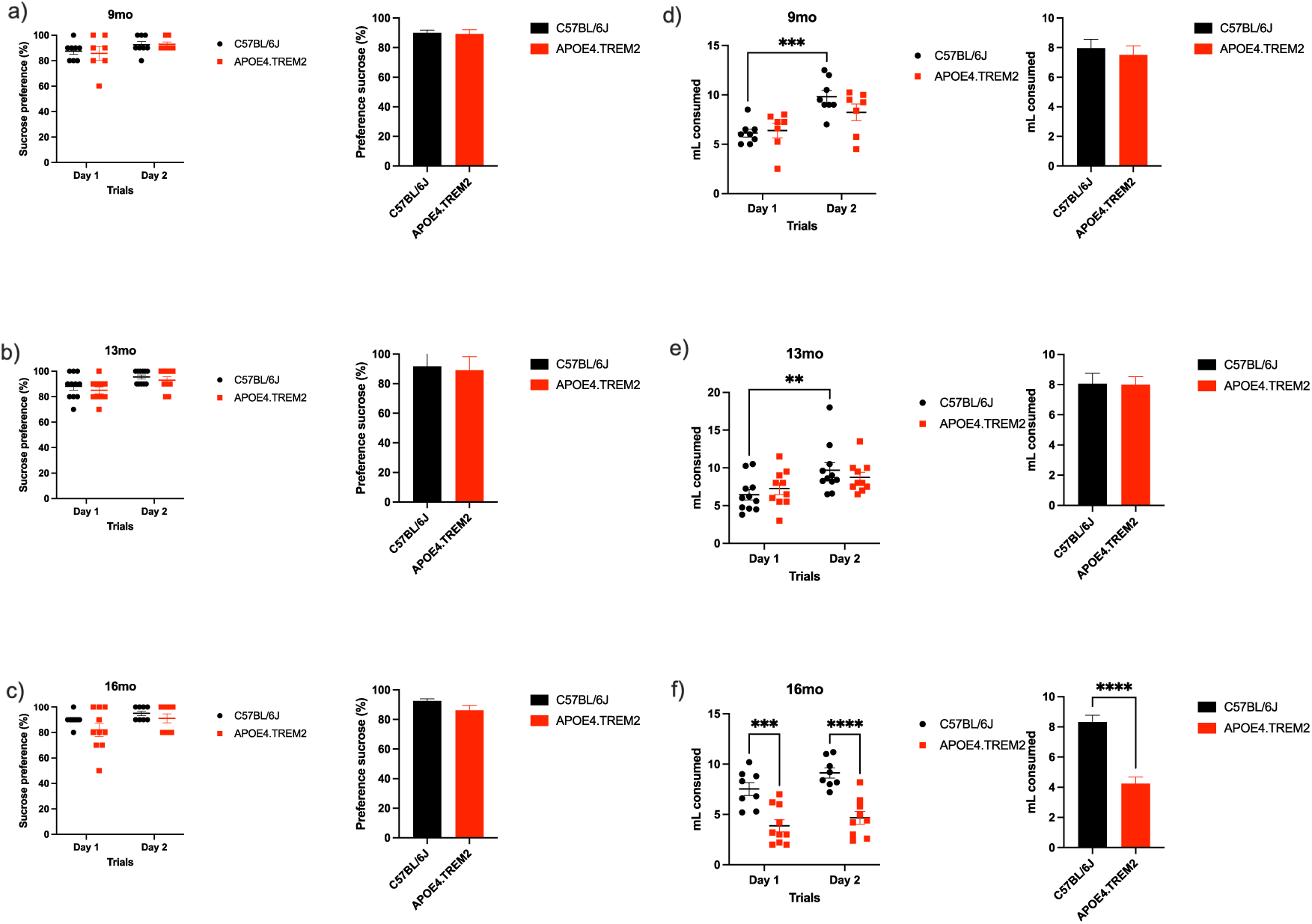
Sucrose Preference Task. Data shows mean ± SEM for the for WT and APOE4.TREM2 preference for sucrose when presented with both regular water and a 10% sucrose solution as a measurement for anhedonia. **(A**) dot plot representing individual animal consumption of sucrose (10%) in water and bar chart illustrating the percentage of sucrose preference for both trials for WT (black) and APOE4.TREM2 (red) at 9 mo, (**B**) for 13 mo WT (black) APOE4.TREM2 (red), and (**C**) 16 mo WT (black) APOE4.TREM2 (red). (**D**) dot plot representing the individual animal consumption (mL) of sucrose in water and the water control, bar chart illustrating the average of consumption over two experimental days for the group at 9 mo, (**E**) at 13 mo, and (**F**) at 16 mo. Significance determined by two-way ANOVA followed by Bonferroni’s post-hoc, *P < 0.05, **P < 0.01, ***P < 0.001, ****P < 0.0001. See **Table 2** for number of mice per group.

Similarly, at 13 mo, both groups showed high preference on the first day, WT 88.2 ± 3.0 % and APOE4.TREM2 85.0 ± 2.7 %. Preference was maintained on the second day for both groups WT 95.5 ± 1.6 % and APOE4.TREM2 93 ± 2.6 % with no statistical differences between groups (F_(1,38)_=1.27, P=0.27; two-way ANOVA, **Figure 2b**).

At 16 mo both groups again showed similar preference with no statistical difference with WT ingesting 90.0 ± 1.9 % and APOE4.TREM2 82.0 ± 5.1 % on the first day. Both groups maintained preference as WT consumed 95.0 ± 1.9 % and APOE4.TREM2 91.1 ± 3.5 % with no statistical difference (F_(1,31)_=2.64, P=0.12; two-way ANOVA, **Figure 2c**).

In addition to preference, the total amount of volume consumed was measured every day. Over the test days, at 9 mo the groups showed a time effect (F_(1,26)_=17.8, P<0.001; two-way ANOVA, **Figure 2d**), and the WT mice at 9 mo increased their consumption from the first day: 6.1 ± 0.4 mL to the second day at 9.8 ± 0.6 mL (P=0.012). Meanwhile, the APOE4.TREM2 consumed 6.4 ± 0.7 mL on the first day and 8.2 ± 0.8 mL on the second day with no statistical differences (P=0.39).

The WT at 13 mo also showed an effect over days (F_(1, 38)_ = 8.97, P = 0.005; two-way ANOVA, **Figure 2e**), with an increase in consumption between the first day 6.4 ± 0.7 mL and the second day of test 9.7 ± 1.0 mL (P = 0.031). APOE4.TREM2 mice did not show statistical differences between first day (7.3 ± 0.8) and the second day (8.8 ± 0.7).

Interestingly, at 16 mo there was only a marginal effect across days (F_(1,31)_=3.87, P=0.059), while there was a strong effect between groups (F_(1,31)_=44.63, P<0.0001; two-way ANOVA, **Figure 2f)**. WT group consumed more on the first day (7.5 ± 0.6 mL) than the APOE4.TREM2 (3.9 ± 0.6 mL, P<0.001). This behavior was observed on the second testing day as the WT consumed 9.1 ± 0.5 mL while the APOE4.TREM2 consumed 4.7 ± 0.6 mL (P<0.0001). Thus, APOE4.TREM2 mice at 16 mo consumed significantly less compared to control. Overall, however, we did not find anhedonia in APOE4.TREM2 mice.

### Open field test: APOE4.TREM2 at 16 months old showed increased anxiety

Anxiety and exploratory activities were evaluated by allowing mice to freely explore an open field arena for 15 minutes in a classic open field-testing apparatus (i.e., a white PVC square arena, 50 x 50 cm, with walls 40 cm high). The mice showed an aging effect (F_(2,50)_ = 8.62, P = 0.0006; two-way ANOVA, **Figure 3a**). Specifically, time spent in the central 50% of the total area was lower in APOE4.TREM2 mice at 16 mo (124.9 ± 12.0 sec) compared to the 9 mo WT (216.1 ± 23.4 sec, P = 0.0096), 9 mo APOE4.TREM2 (213.5 ± 16.7 sec, P = 0.014), and 13 mo WT (225.1 ± 16.0 sec, P = 0.001) (**Figure 3a**). Genotype was marginally significant (F_(2,50)_=3.49, P = = 0.068).

**Figure 3.**
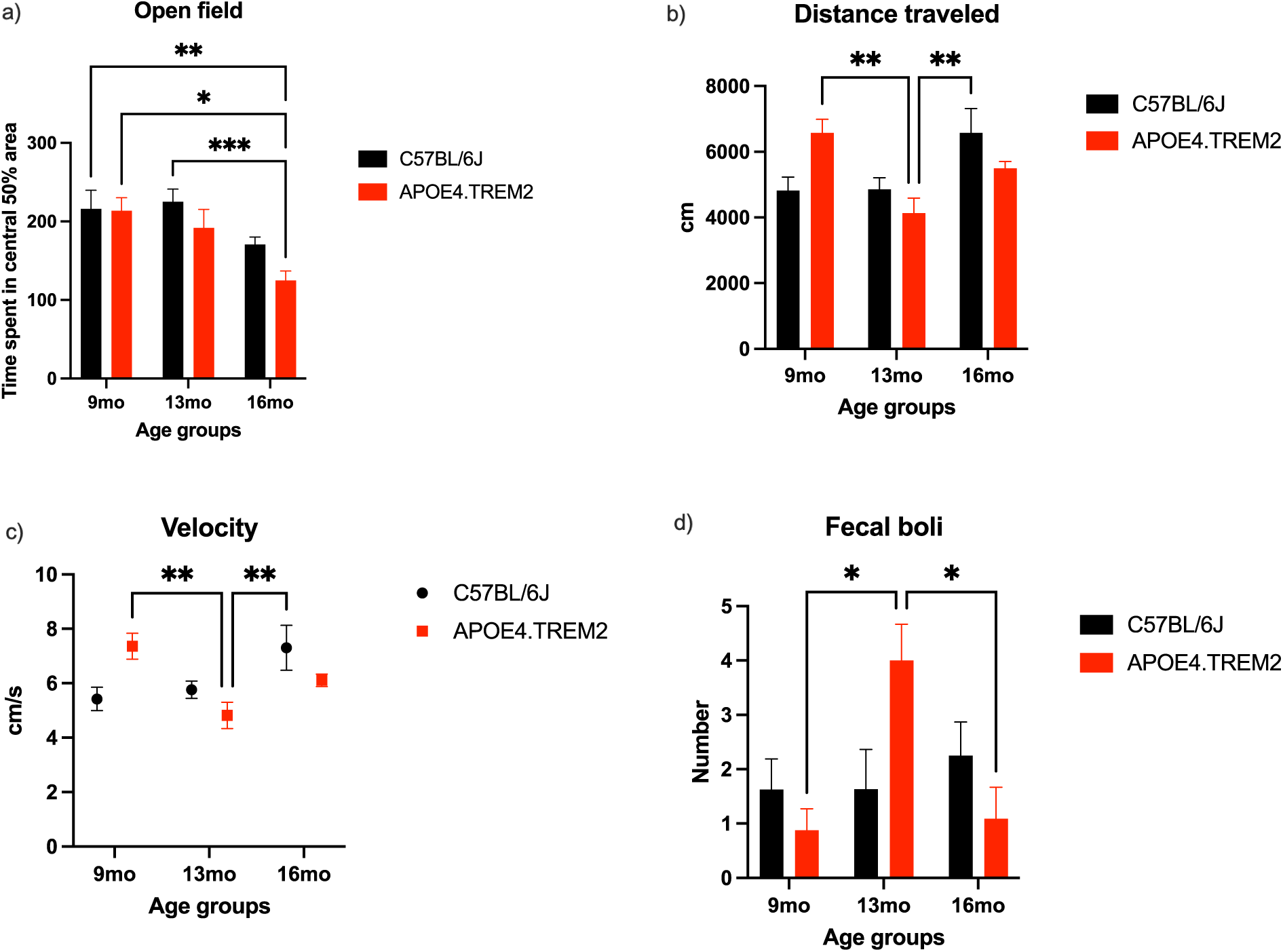
Open Field Test. Bar charts with mean ± SEM of time mice spent in central 50% of arena (**A**) and their distance traveled (mean ± SEM (cm; **B**). Velocity averaged across age and genotype with mean ± SEM (cm/s; **C**). Bar chart showing fecal production (**D**) with mean ± SEM for WT (black) and APOE4.TREM2 (red) mice. Significance determined by two-way ANOVA followed by Bonferroni’s post-hoc, *P < 0.05, **P < 0.01, ***P < 0.001. See **Table 2** for number of mice per group.

During these trials the total distance traveled (cm), and velocity (cm/s) were also measured (**Figures 3b and 3c**), with a significant effect of age (F_(2,50)_ = 7.51, P=0.002) and an age:genotype interaction (F_(2,50)_ = 5.84, P=0.006; two-way ANOVA; **Figure 3b**) for distance, and hence similarly for velocity (age (F_(2,50)_ = 5.51, P=0.007) and age:genotype interaction (F_(2,50)_ = 6.51, P=0.004; two-way ANOVA), **Figure 3c**). The APOE4.TREM2 mice at 13 mo traveled (4138 ± 454 cm), less than the APOE4.TREM2 mice at 9 mo (6578 ± 409 cm; P=0.005) and to WT at 16 mo (6576 ± 742 cm; P=0.005). The 13 mo APOE4.TREM2 showed lower velocity (4.8 ± 0.5 cm/sec) compared to APOE.TREM2 at 9 mo (7.4 ± 0.5 cm/s, P = 0.006) and WT 16 mo (7.3 ± 0.8 cm/sec, P = 0.008).

Additionally, the count of fecal boli was obtained at the end of the test and showed main effect of genotype (F_(2,50)_=3.41, P = 0.042) on the number of fecal boli and an age:genotype interaction (F_(2,50)_ = 4.98, P = 0.011; two-way ANOVA, **Figure 3d**). At 13 mo, APOE4.TREM2 mice had a higher count of fecal boli (4.0 ± 0.7) compared to 9 mo (0.9 ± 0.4, P = 0.018) and compared to 16 mo (1.1 ± 0.6, P=0.016). These findings demonstrate age dependent increase in anxiety in APOE4.TREM2 mice.

### Hidden Cookie Test: APOE4.TREM2 at 13 mo showed deficits in the Cookie Test

A quarter of chocolate cookie was placed every 48 hours three times in the home cage on the top of the bedding. After the familiarization phase, mice were deprived of the cookie for 3 days before the test. On the first day of the test, a piece of cookie was buried in one corner of their home cage and the latency to finding the cookie was recorded. An hour and a half later, another piece of cookie was buried in the same corner as the first trial. The next day, mice were placed in the experimental cages and the piece of cookie was buried in a corner different from the prior day. This was tested twice following the same protocol as the first experimental day.

Mice did not show statistical differences between genotypes (F_(1,48)_ = 0.17, P = 0.68; two-way ANOVA, **Fig. 4a**) at 9 mo or at 16 mo (F_(1,64)_ = 0.45, P = 0.50; two-way ANOVA, **Fig. 4c**). Statistical significance was seen in 13 mo APOE4.TREM2 mice, as they showed greater latency in finding the cookie compared to WT, specifically on the last trials of each day (F_(1,76)_ = 21.92, P < 0.0001; two-way ANOVA, **Fig. 4b**).

**Figure 4.**
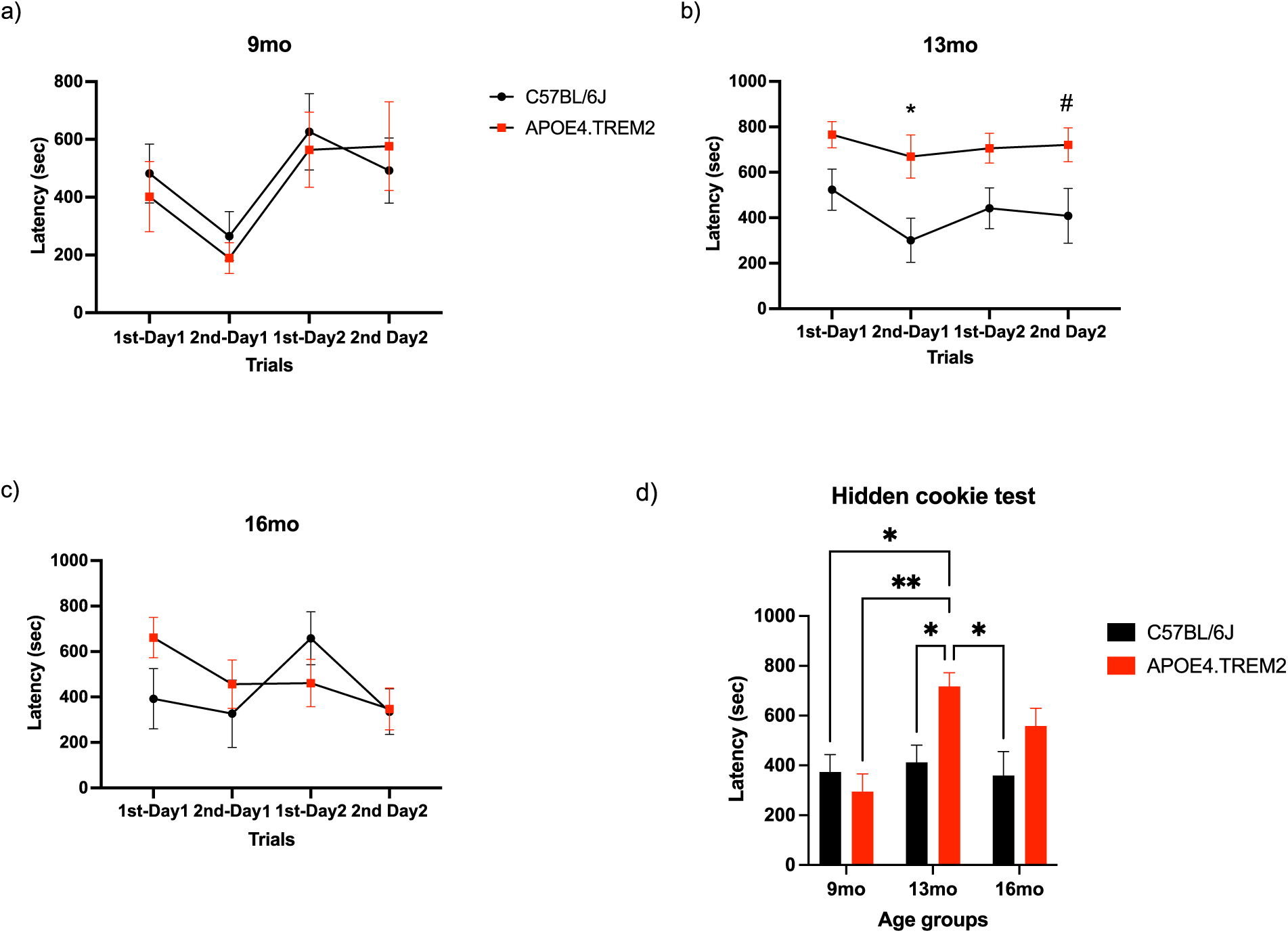
Hidden Cookie Test. Data depicts mean ± SEM for latency to cookie detection for both days at (**A**) 9 mo old WT (black) and APOE4.TREM2 (red), (**B**) 13 mo old WT (black) and APOE4.TREM2 (red), and (**C**) 16 mo old WT (black) APOE4.TREM2 (red). (**D**) Bar graph represents the mean ± SEM for latency of trials on day 1 by age and genotype. ^#^P=0.07, *P < 0.05, **P < 0.01.

Figure 4d compares the latencies averaged over the experimental days on day 1, showing significant effect of age (F_(2,100)_ = 4.02, P = 0.011), genotype (F_(1,100)_=5.52, P=0.021) and age: genotype interaction (F_(2, 100)_ = 3.37, P = 0.04; two-way ANOVA). The APOE4.TREM2 at 13 mo showed higher latency (717.5 ± 55 sec) compared to 13 mo WT (412.5 ± 69 sec; P = 0.023). Additionally, 13 mo APOE4.TREM2 mice showed differences compared to WT at 9 mo (373.6 ± 69.7 sec, P = 0.016), to 9 mo APOE4.TREM2 (295.4 ± 70.9 sec, P = 0.004), and 16 mo WT (359.5 ± 95.8 sec, P = 0.016). The increased latency in buried cookie detection, especially in 13 mo APOE4.TREM2 mice, indicates genotype related dysfunction of olfactory guided foraging tasks.

### Morris Water Maze: APOE4.TREM2 show deficits in the MWM Task

The MWM assesses spatial memory and learning deficits allowing mice to swim freely in a pool, first learning the location of the escape platform. On the test day, the platform was removed, and each mouse was allocated 120 sec to swim freely. During this time, the latency to the platform area, cumulative time at the platform area, and cumulative time in the quadrant of the platform area were measured.

Latency was only affected by genotype (F_(1,42)_ = 7.16, P <0.011; two-way ANOVA; Fig. 5a). At 9 and 16 mo there were no differences. Mice at 13 mo showed a higher latency (49.4 ± 17.6 sec) to find the platform area compared to WT mice (14.5 ± 3.2 sec; P = 0.011).

**Figure 5.**
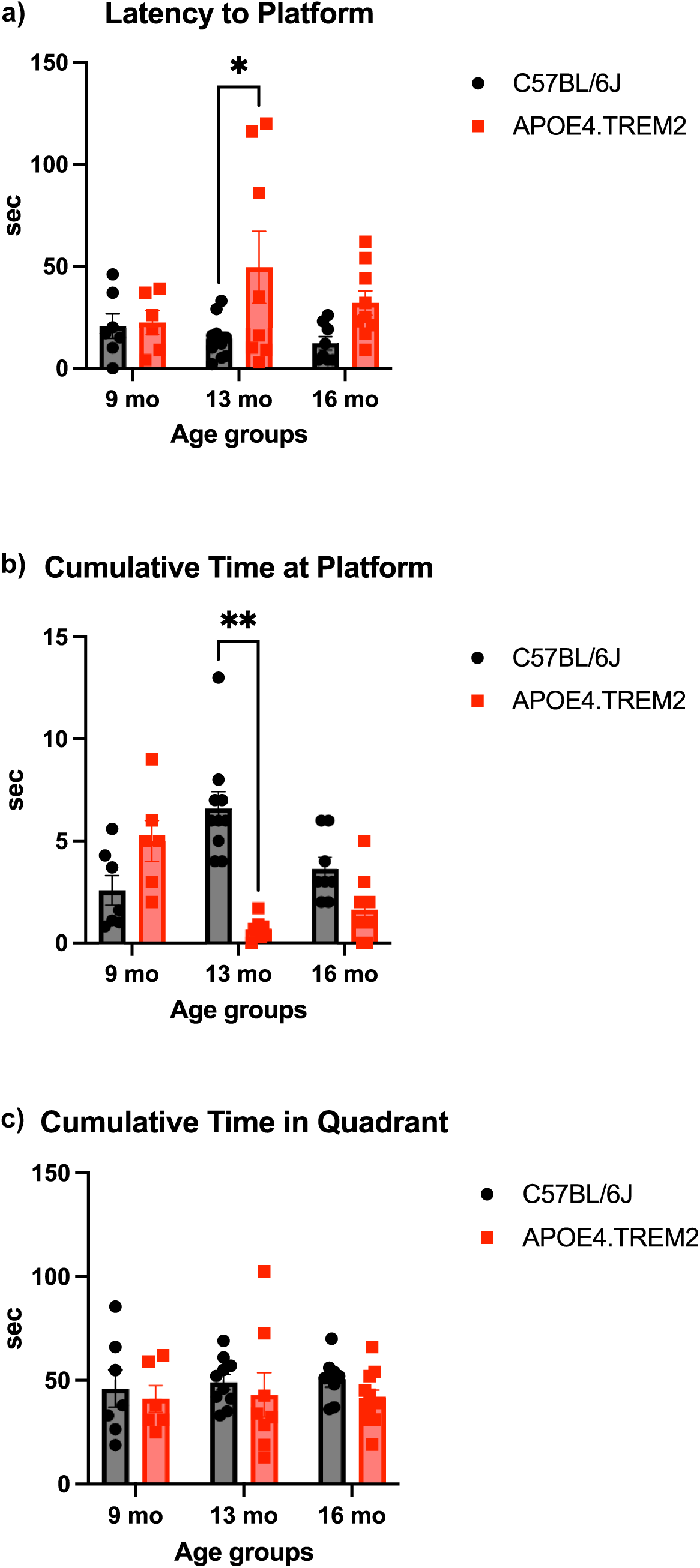
Morris Water Maze. Data displays WT (black) and APOE4.TREM2 mice (red) by age. (**A**) Bar graph representation of mean ± SEM latency to platform area. (**B**) Bar graph displays the mean ± SEM cumulative time spent in platform area. (**C**) Bar graph of cumulative time spent in quadrant. There were no statistical differences for the cumulative time in the quadrant. Significance determined by two-way ANOVA followed by Bonferroni’s post-hoc, *P < 0.05, **P < 0.0001.

Time at the platform was affected by both genotype (F_(1,44)_ = 11.61, P <0.002) and age:genotype interaction (F_(2,44)_ = 18.81, P <0.0001; two-way ANOVA; Fig. 5b). The 13 mo APOE4.TREM2 mice spent significantly less time at the platform area (0.7 ± 0.2 sec) compared to the WT (6.6 ± 0.8 sec; P<0.0001).

There were no statistical differences for the cumulative time in the quadrant (Fig. 5c). We can conclude from these results that the APOE4.TREM2 mice display genotype-dependent deficits in spatial memory and learning.

### Nestlet building: APOE4.TREM2 do not show deficits in nest building

The nestlet building test measures the capacity of mice to build a nest to evaluate innate behavior, motor deficits, as well as overall welfare. Mice are scored out of 5 points, 1 indicates a >90% intact nestlet, whereas a 5 indicates a nestlet torn >90% and a clear nest crater [34]. Scores were averaged across three raters using anonymized nest pictures.

Two-way ANOVA showed only an interaction effect, of age and genotype (F_(2,46)_ = 4.51, P = 0.017; Figure 6). At 9 mo both WT and APOE4.TREM2 mice scored similarly (3.4 ± 0.1 vs 3.5 ± 0.3) without statistical difference. At 13 mo, the WT scored 4.3 ± 0.3, higher than APOE4.TREM2 at 3.2 ± 0.3 (P = 0.005). At 16 mo, the control group scored 3.1 ± 0.4, similar to the APOE4.TREM2 3.5 ± 0.1. WT mice scored higher at 16 mo than 13 mo (P = 0.006). Overall, no impairment in nestlet building in APOE4.TREM2 mice is evident.

**Figure 6.**
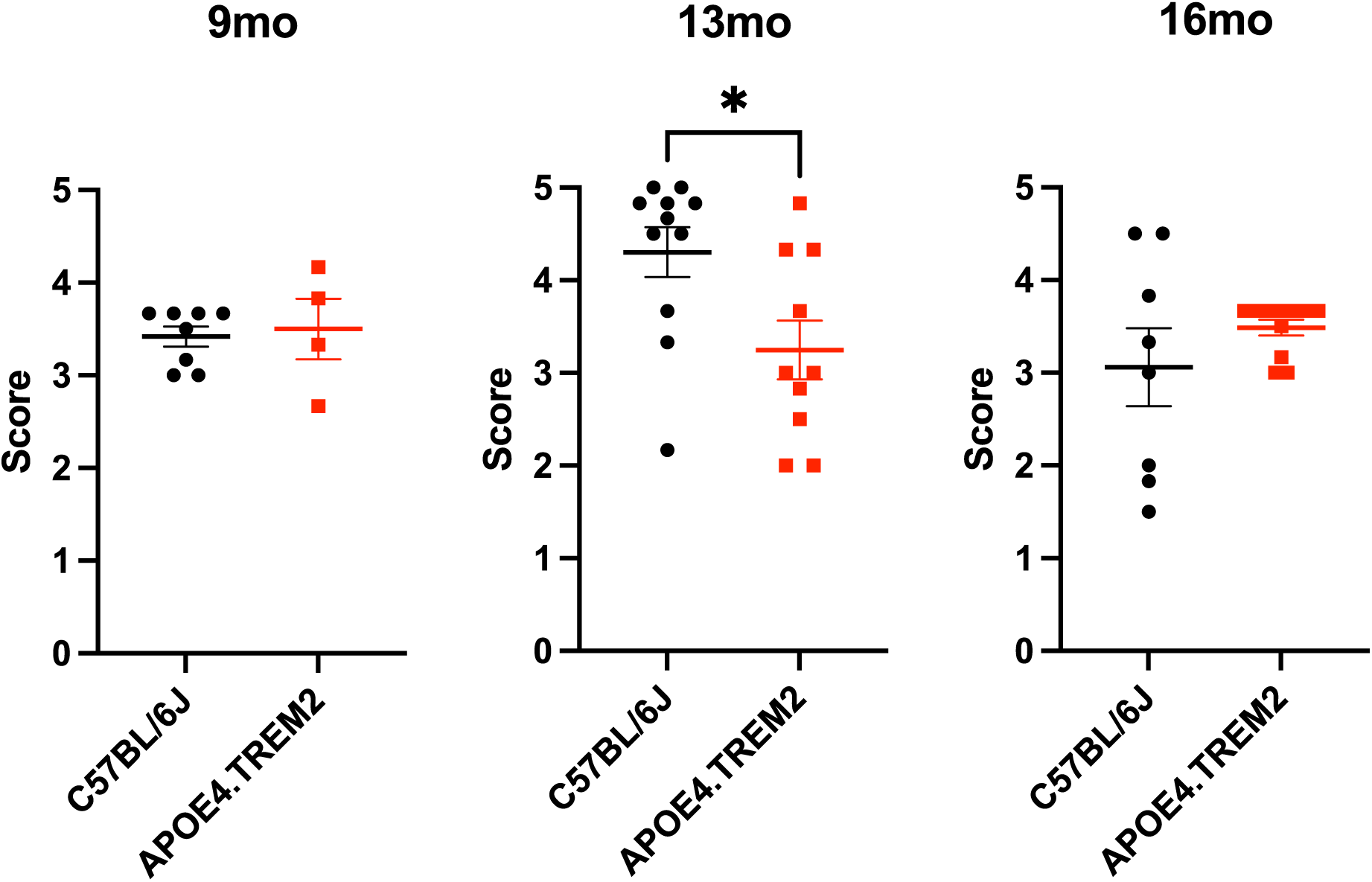
Nestlet building. Dot plots with mean ± SEM of nestlet building scores for WT (black) and APOE4.TREM2 (red) out of 5 points, 1 indicates a >90% intact nestlet, whereas a 5 indicates a nestlet torn >90% and a clear nest crater [34] at 9 mo (**A**), 13 mo (**B**), and 16 mo (**C**). *P < 0.01.

### Sleep/wake cycle: APOE4.TREM2 did not show disturbances in the sleep/wake cycle

The sleep/wake cycle was measured for two days to know if the APOE4.TREM2 mice showed sleep alterations. At 9 mo, both groups were active when they were in the dark phase of the experiment (Fig. 7a, WT Day 1: 18.5 ± 3.3 % of dark time was slept and Day 2: 29.9 ± 4.4 %; APOE4.TREM2 Day1: 23.3 ± 4.9 % and Day 2: 32.4 ± 2.7 %). Meanwhile, the percentage of sleep was higher during the light phase than dark phase for both groups. (Fig. 7a, WT Day 1: 56.8 ± 1.8 % and Day 2: 54.1 ± 1.4 %; APOE4.TREM2 Day 1: 57.6 ± 1.5 % and Day 2: 54.6 ± 1.0 %; both post-hoc P < 0.001) (phase effect F_(3,52)_= 70.70, P<0.0001; two-way ANOVA). Differences were also not found between groups when both days were averaged (Fig. 7d, dark phase WT 24.2 ± 3.0 % vs APOE4.TREM2 27.9 ± 3.0 %; light phase WT 55.4 ± 1.2 % vs APOE4.TREM2 56.1 ± 1.0 %) (F_(1,56)_= 0.90, P=0.35; two-way ANOVA).

**Figure 7.**
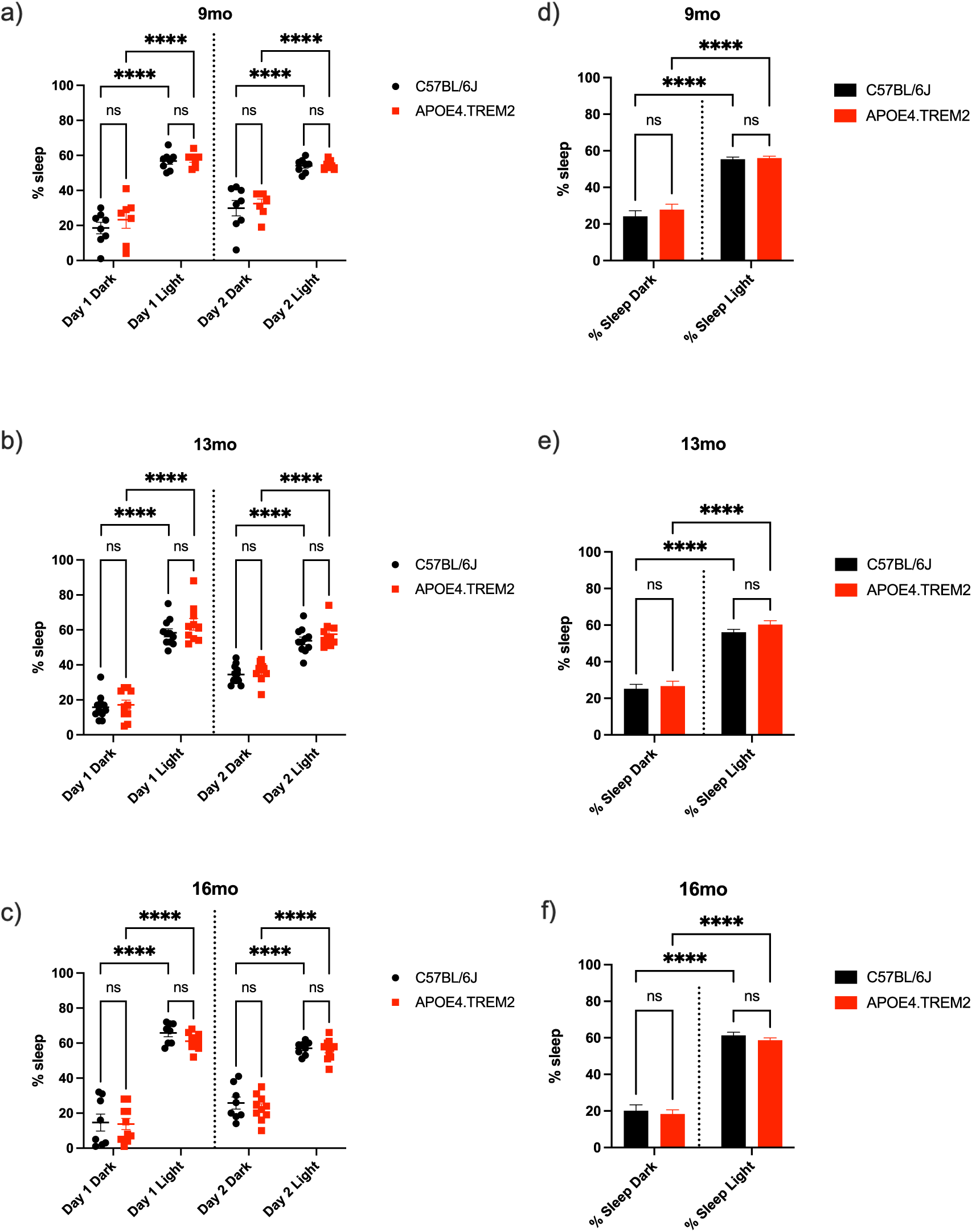
Sleep-wake cycle. Data displays WT (black) and APOE4.TREM2 mice (red) by age. (**A**) Dot plot illustrating the percentage of sleep for individual animals at 9 mo old during both dark and light phases for both days. (**B**) Dot plot illustrating the percentage of sleep for individual animals at 13 mo old during both dark and light phases for both days. (**C**) Dot plot illustrating the percentage of sleep for individual animals at 16 mo old during both dark and light phases for both days. Bar charts comparing average percent sleep over both days in light and dark phase for WT and APOE4.TREM2 at 9 mo (**D**), 13 mo (**E**), and 16 mo (**F**). ****P<0.0001.

At 13 mo, the WT and APOE mice increased their sleep time in the dark phase of the experiment across days (Fig. 7b, WT Day 1: 15.8 ± 2.1 % vs Day 2: 34.6 ± 1.6 %; APOE4.TREM2 Day 1: 17.2 ± 2.7 % vs Day 2: 36.2 ± 1.8 %, both post-hoc P <0.0001), but their higher (both P<0.0001) % sleep in the light phase didn’t differ across days (Fig. 7b, WT Day 1: 58.4 ± 2.3 % vs Day 2: 53.8 ± 2.2 %; APOE4.TREM2 Day 1: 63.2 ± 3.4 % vs Day 2: 57.5 ± 2.3 %) (phase effect F_(3,76)_= 153.1, P<0.0001; two-way ANOVA). Data did not show statistical difference for the total sleep between groups when both days were averaged (Fig. 7e, dark phase WT 25.2 ± 2.4 % vs APOE4.TREM2 26.7 ± 2.7 %; light phase WT 56.1 ± 1.6 % vs APOE4.TREM2 60.4 ± 2.1 %) (F_(1,80)_= 0.1.69, P=0.20; two-way ANOVA).

At 16 mo, both groups were also most active in the dark phase (Fig. 7c, WT Day 1: 14.6 ± 4.8 % vs Day 2: 25.8 ± 3.5 %; APOE4.TREM2 Day 1: 13.8 ± 3.2 % vs Day 2: 23.0 ± 2.3 %).

Both groups increased their sleep percentage in the light phase (Fig. 7c, WT Day1: 65.8 ± 2.1 % and Day 2: 57.0 ± 1.3 %; APOE4.TREM2 Day1: 61.1 ± 1.5 % and Day 2: 56.3 ± 1.9 %; both P<0.0001) (phase effect F_(3,64)_= 155.1, P<0.0001; two-way ANOVA). Again, no statistical difference between groups was observed for the total percentage of sleep in the two-day average of the dark/light cycle (Fig. 7f, dark phase WT 20.2 ± 3.2 % vs APOE4.TREM2 18.4 ± 2.2 %; light phase WT 61.4 ± 1.6 % vs APOE4.TREM2 58.7 ± 1.3 %) (F_(1,68)_= 1.08, P=0.31; two-way ANOVA).

We conclude that we did not find age or genotype related differences in sleep time across the sleep/wake cycle.

### Odor-evoked intrinsic response signaling: vascular hyperresponsiveness shows genotype and age dependent effects in the APOE4.TREM2 model

Odor-evoked intrinsic signals were recorded and measured in anesthetized ∼0.5 to ∼2 years old mice (**Table 3**) to evaluate whether the APOE4.TREM2 genotype produces vascular hyperresponsiveness, particularly with increasing age. The responses were calculated for 3 types of region of interest (ROI): 1SD above baseline, 3SD above baseline and un-thresholded (Fig. 8A**-C** and **Methods**). The 3SD ROI type reflects dOB arteries, as verified by FITC injection (Fig. 8E, Methods). Arteriolar vessels cannot be selected based on resting light image vessels alone (Fig. 8D), which can include non-arterial vessels.

**Figure 8.**
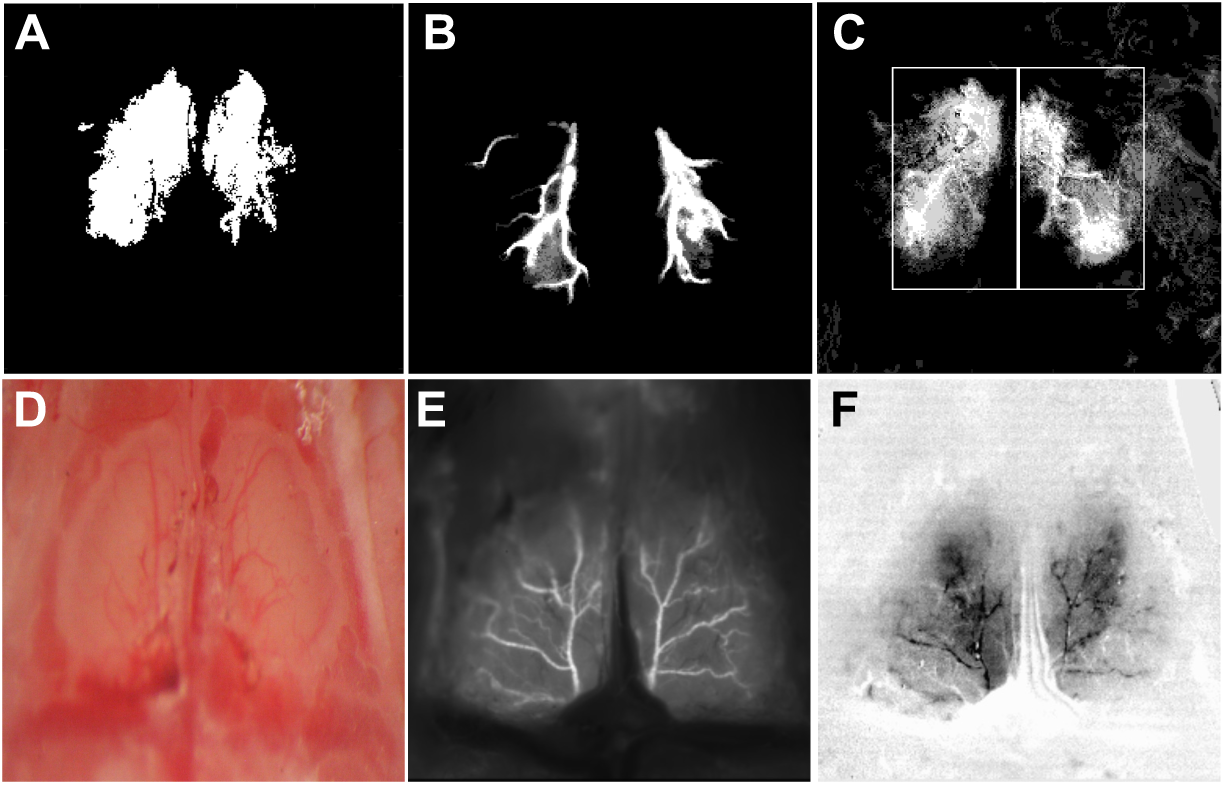
ROIs and dOB. To ascertain robustness of our findings we defined 3 types of ROI. (A) ROI 1 SD: dOB pixels with a blood volume (BV) change of 1 SD > baseline, in 4/6 trials. (B) ROI 3 SD: 3 SD > baseline in 3/6 trials. (C) ROI Rectangle: anatomically based ROI analyzing two rectangular areas (L dOB and R dOB) with no thresholds. (D) dOB under ambient light post-surgery. (E) dOB under 488nm LED with FITC injection, resembling the 3SD ROI. (F) dOB under 590nm LED (see Methods).

The dOB was imaged using a 590nm LED, the power spectrum of which matches hemoglobin’s isosbestic point and therefore reports changes in blood volume (BV). An increase in BV increases absorption and hence decreases reflected signal. The average hemodynamic response function shown in Fig. 9 represents BV dOB responses to odor presentations (flow dilutions at 0%, 0.3%, 1%, and 3% of saturated vapor pressure) of ethyl butyrate (EB) for 4 sec and was calculated based on the 3SD-based ROI across all mice (n=38, **Table 3**) and trials (n=12/mouse for EB). These responses have a rapid (∼1 sec) onset and a slow offset lasting over 10 sec and have larger amplitude with increasing odor concentrations. The data demonstrate that the APOE4.TREM2 mice show increased intrinsic BV responses (i.e. hyperresponsiveness) at each concentration of EB, being especially prominent at 1 and 3% EB, relative to WT.

**Figure 9.**
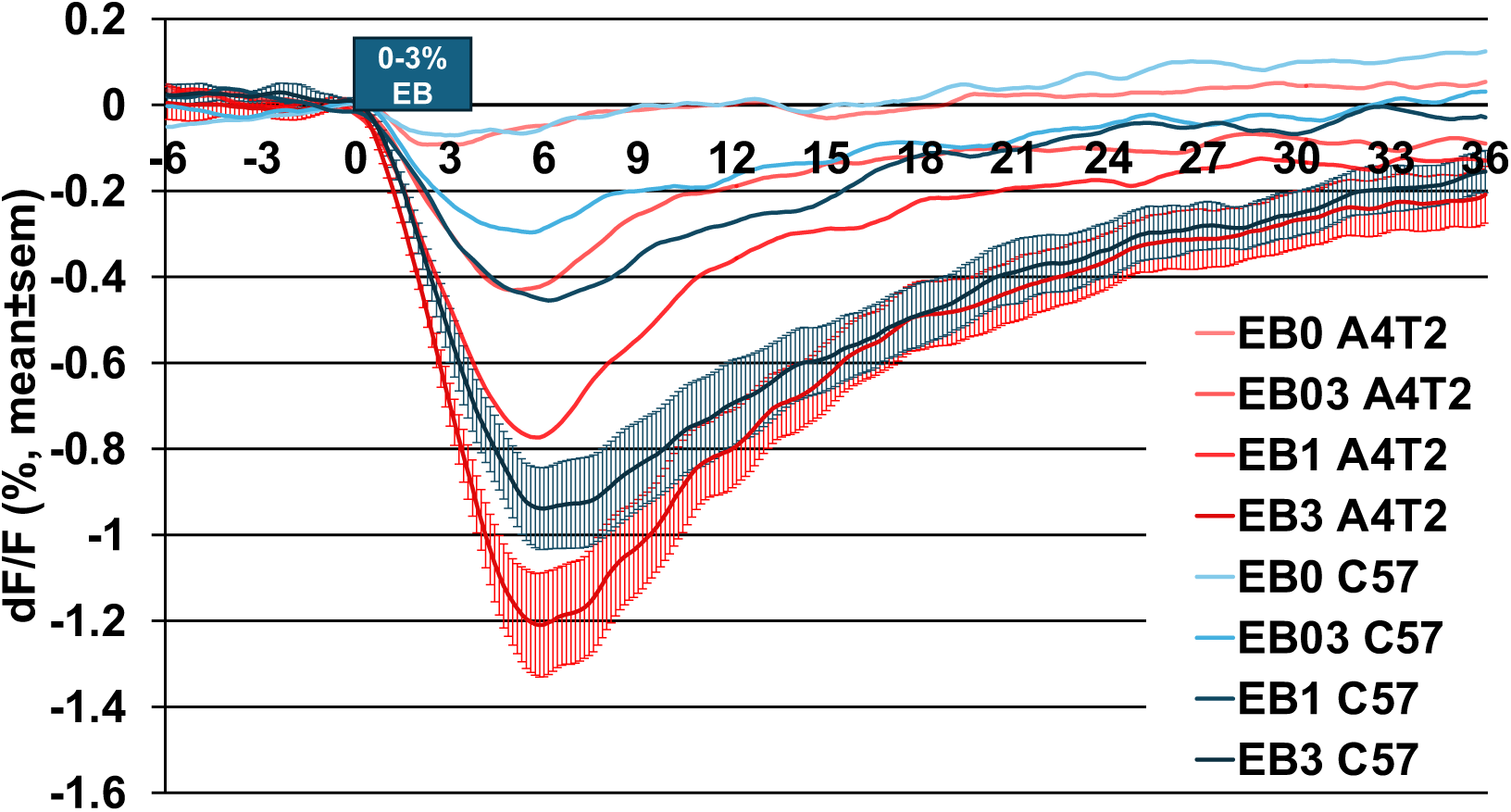
Hemodynamic response function. Averaged odor-evoked intrinsic BV response comparison of the two genotypes at each concentration of 4s of EB (0% control, 0.3%, 1.0%, 3.0%). WT (black) and APOE4.TREM2 (red). Intrinsic hyperresponsiveness is seen in APOE4.TREM2 at each concentration. See **Table 3** for number of mice per group.

We calculated the Area Under the Curve (AUC) to assess the overall integrated BV response magnitude over the first 10s post odor onset (sum of % dF/F across 70 frames, i.e. %dF/F * t (10 sec) * frame rate (7 frames per sec)). A more negative AUC value signifies greater BV response. Evaluating the AUC ignores the response dynamics to focus on the overall magnitude. When AUC is averaged across mice combining both odorants in each ROI, APOE4.TREM2 mice display vascular hyperresponsiveness to odor stimuli at each concentration. Statistically highly significant odor-induced BV hyperresponsiveness by APOE4.TREM2 mice is evident in the 1SD-based ROI responses (Fig. 10a), artery-like 3SD ROI responses (Fig. 10b) and even dOB-wide rectangular ROI responses (Fig. 10c). **Table 4 and 5** demonstrate, using 2-way (age, genotype) and 3-way (age, genotype, sex) ANOVA across all mice and all 24 trials per mouse, that age and genotype separately and in interaction significantly affect intrinsic responses. Sex was not a significant factor, although we note uneven distribution of males vs. females (**Table 3**). These data demonstrate that the intrinsic odor responses are larger in the APOE4.TREM2 mice, and more so with age.

**Figure 10.**
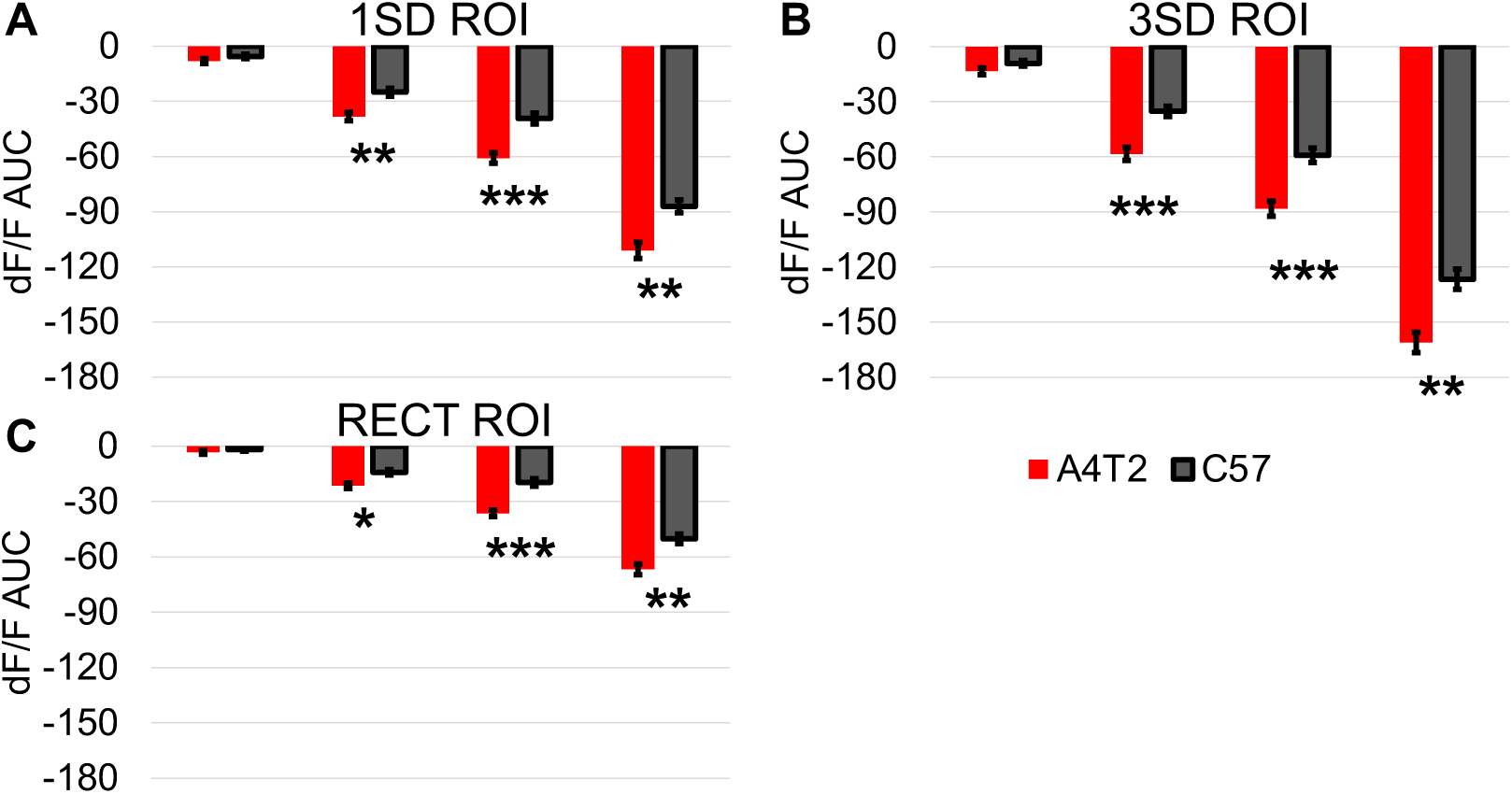
Averaged intrinsic responses comparing genotype at each concentration for all ROIs. Bar charts displaying averaged measurements of the area under the curve (AUC; dF/F(%)*10 s*7fps) combining EB and MV with 3 repetitions per concentration, comparing WT (black) and APOE4.TREM2 (red). A more negative value in the curve signifies greater blood volume changes. (**A**) APOE4.TREM2 vs. WT BV comparison for 1SD threshold-based ROI for 0.3% (P=0.01), 1.0% (P=0.002), and 3% (P=0.006). (**B**) APOE4.TREM2 vs. WT BV comparison for 3SD threshold-based ROI for 0.3% (P=0.003), 1.0% (P=0.003), and 3% (P=0.006). (**C**) APOE4.TREM2 vs. WT BV comparison for the rectangular unthresholded ROI for 0.3% (P=0.03), 1.0% (P=0.004), and 3% (P=0.007). All tests are planned unpaired one-tailed Student’s t-test. *P < 0.05, **P < 0.01, ***P<0.005.

**Table 4.**
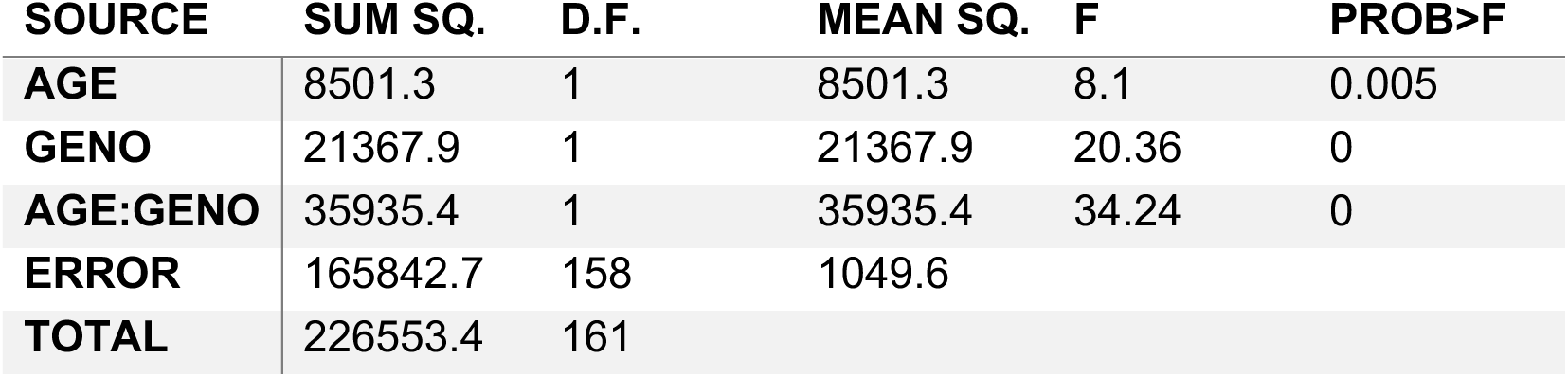
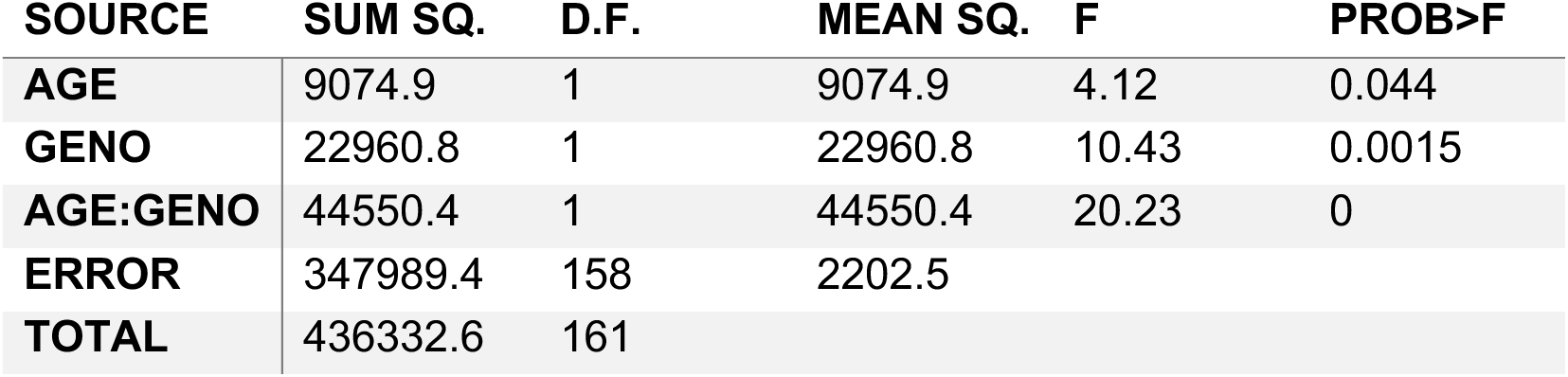
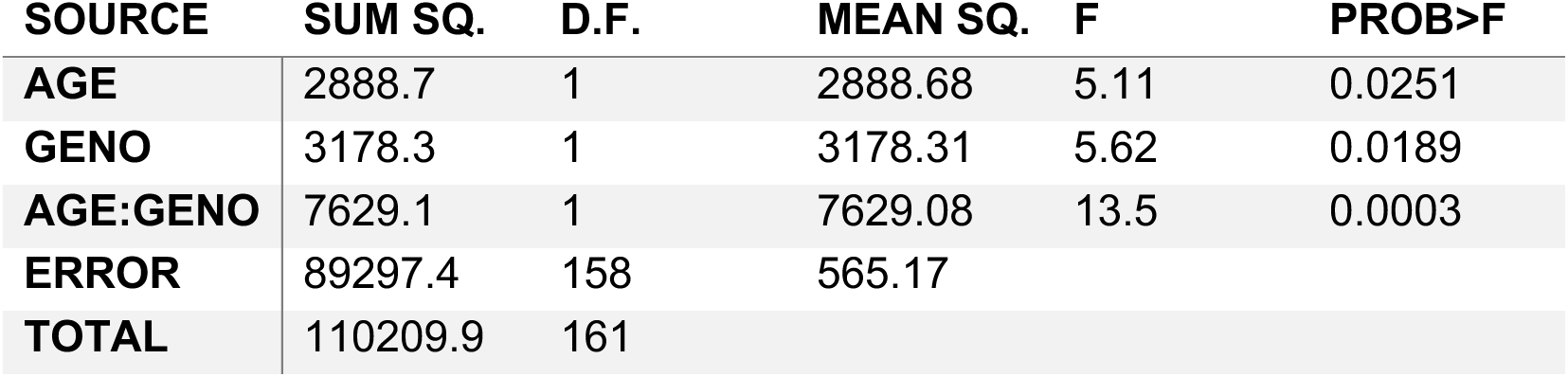
2-way ANOVA confirming genotype and age effects on neurovascular hyperresponsiveness in each RIO (3% EB/MV. ) Statistical differences in the data support odor-evoked intrinsic responses are age and genotype dependent. (**A**) 1 SD ROI type age effect, genotype effect, and age:genotype interaction effect. (**B**) 3 SD ROI type age effect, genotype effect, and age:genotype effect. (**C**) Rectangular ROI type age effect, genotype effect, and age:genotype effect.

| **SOURCE** | **SUM SQ.** | **D.F.** | **MEAN SQ.** | **F** | **PROB>F** |
| --- | --- | --- | --- | --- | --- |
| **AGE** | 8501.3 | 1 | 8501.3 | 8.1 | 0.005 |
| **GENO** | 21367.9 | 1 | 21367.9 | 20.36 | 0 |
| **AGE:GENO** | 35935.4 | 1 | 35935.4 | 34.24 | 0 |
| **ERROR** | 165842.7 | 158 | 1049.6 |  |  |
| **TOTAL** | 226553.4 | 161 |  |  |  |
| **AGE** | 9074.9 | 1 | 9074.9 | 4.12 | 0.044 |
| **GENO** | 22960.8 | 1 | 22960.8 | 10.43 | 0.0015 |
| **AGE:GENO** | 44550.4 | 1 | 44550.4 | 20.23 | 0 |
| **ERROR** | 347989.4 | 158 | 2202.5 |  |  |
| **TOTAL** | 436332.6 | 161 |  |  |  |
| **AGE** | 2888.7 | 1 | 2888.68 | 5.11 | 0.0251 |
| **GENO** | 3178.3 | 1 | 3178.31 | 5.62 | 0.0189 |
| **AGE:GENO** | 7629.1 | 1 | 7629.08 | 13.5 | 0.0003 |
| **ERROR** | 89297.4 | 158 | 565.17 |  |  |
| **TOTAL** | 110209.9 | 161 |  |  |  |

**Table 5.**
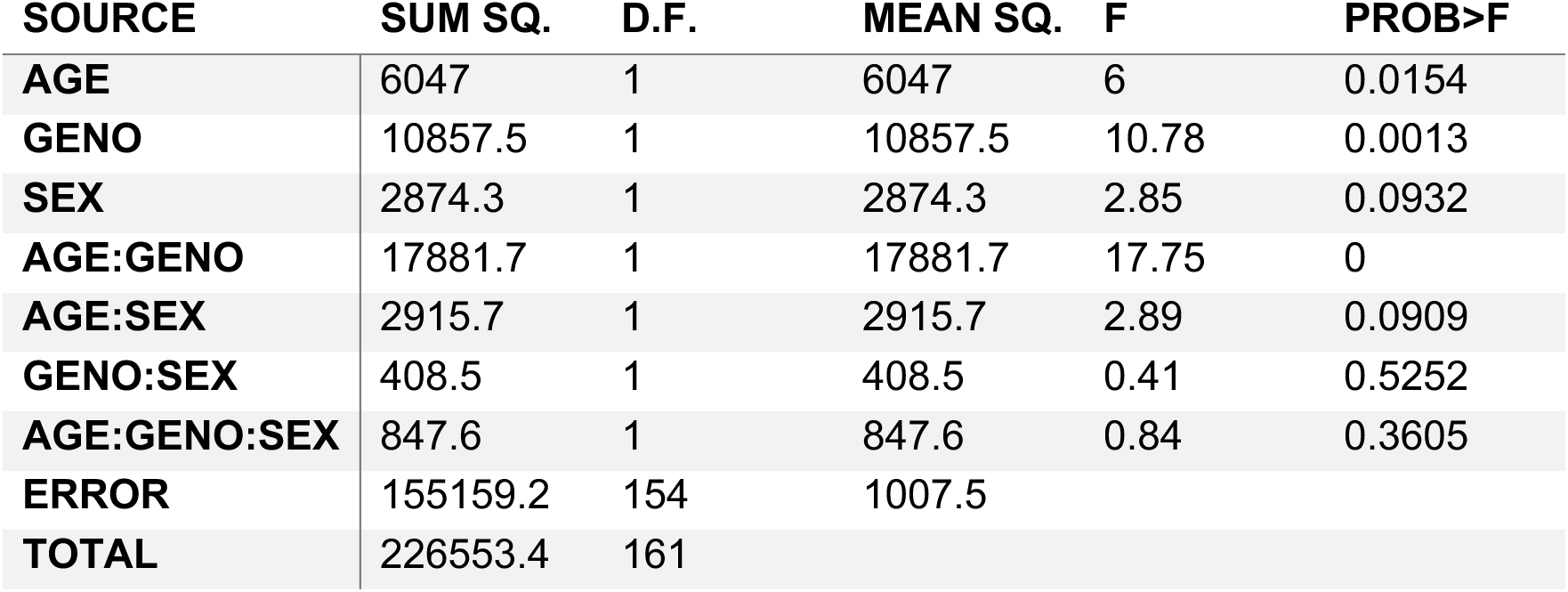
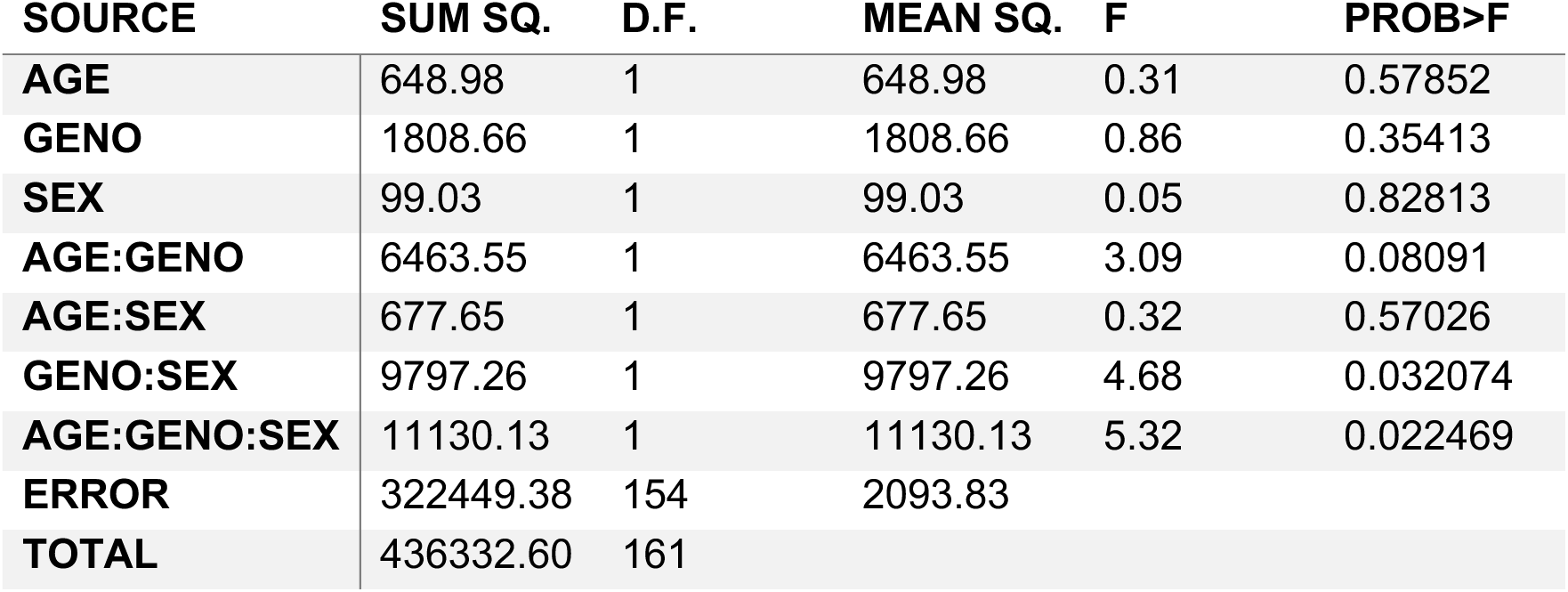
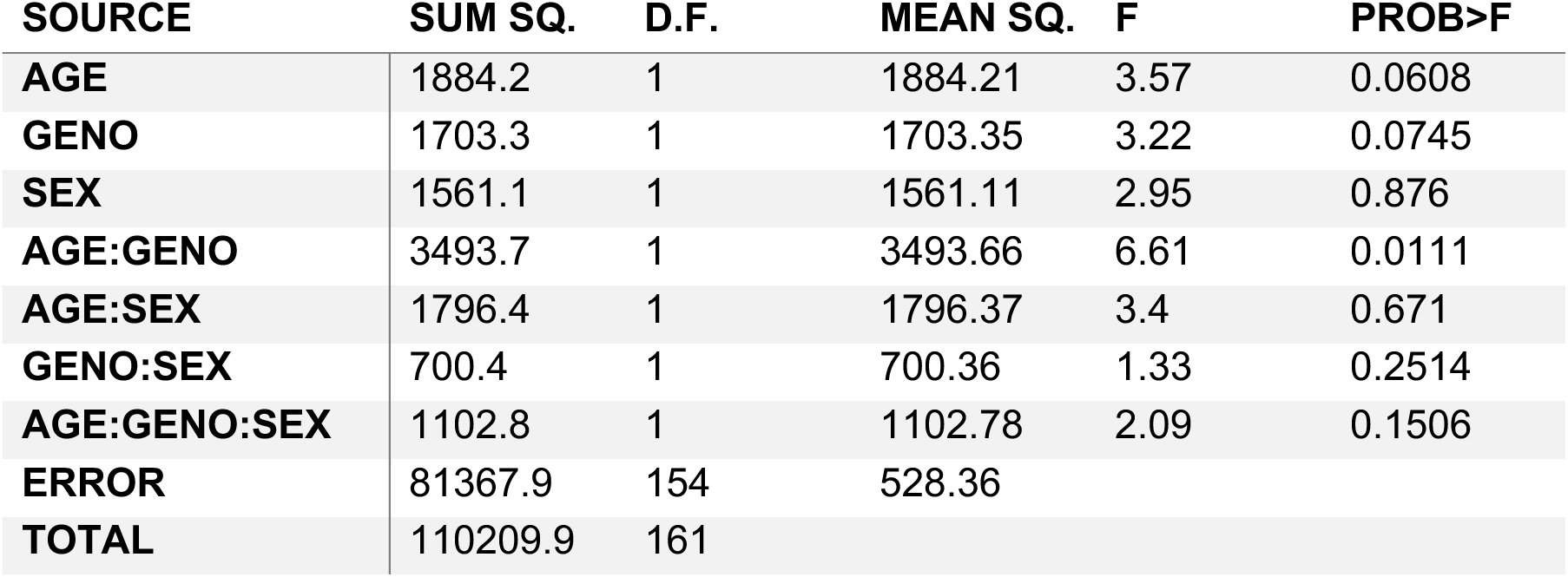
3-way ANOVA assessing genotype, age, and sex effects on intrinsic responses for 3% EB/MV in each ROI. (**A**) 1 SD ROI type responses show significant effect of ge, genotype and their interaction, but no indication of sex related deficits. (**B**) 3 SD ROI type responses show no significant effect of age, genotype, or sex on vascular hyperresponsiveness individually, but show significant interactions. (**C**) Rectangular ROI type data show no significant effect of age, genotype, or sex on vascular hyperresponsiveness individually, but do show a significant genotype:age interaction.

| SOURCE | SUM SQ. | D.F. | MEAN SQ. | F | PROB>F |
| --- | --- | --- | --- | --- | --- |
| AGE | 6047 | 1 | 6047 | 6 | 0.0154 |
| GENO | 10857.5 | 1 | 10857.5 | 10.78 | 0.0013 |
| SEX | 2874.3 | 1 | 2874.3 | 2.85 | 0.0932 |
| AGE:GENO | 17881.7 | 1 | 17881.7 | 17.75 | 0 |
| AGE:SEX | 2915.7 | 1 | 2915.7 | 2.89 | 0.0909 |
| GENO:SEX | 408.5 | 1 | 408.5 | 0.41 | 0.5252 |
| AGE:GENO:SEX | 847.6 | 1 | 847.6 | 0.84 | 0.3605 |
| ERROR | 155159.2 | 154 | 1007.5 |  |  |
| TOTAL | 226553.4 | 161 |  |  |  |
| AGE | 648.98 | 1 | 648.98 | 0.31 | 0.57852 |
| GENO | 1808.66 | 1 | 1808.66 | 0.86 | 0.35413 |
| SEX | 99.03 | 1 | 99.03 | 0.05 | 0.82813 |
| AGE:GENO | 6463.55 | 1 | 6463.55 | 3.09 | 0.08091 |
| AGE:SEX | 677.65 | 1 | 677.65 | 0.32 | 0.57026 |
| GENO:SEX | 9797.26 | 1 | 9797.26 | 4.68 | 0.032074 |
| AGE:GENO:SEX | 11130.13 | 1 | 11130.13 | 5.32 | 0.022469 |
| ERROR | 322449.38 | 154 | 2093.83 |  |  |
| TOTAL | 436332.60 | 161 |  |  |  |
| AGE | 1884.2 | 1 | 1884.21 | 3.57 | 0.0608 |
| GENO | 1703.3 | 1 | 1703.35 | 3.22 | 0.0745 |
| SEX | 1561.1 | 1 | 1561.11 | 2.95 | 0.876 |
| AGE:GENO | 3493.7 | 1 | 3493.66 | 6.61 | 0.0111 |
| AGE:SEX | 1796.4 | 1 | 1796.37 | 3.4 | 0.671 |
| GENO:SEX | 700.4 | 1 | 700.36 | 1.33 | 0.2514 |
| AGE:GENO:SEX | 1102.8 | 1 | 1102.78 | 2.09 | 0.1506 |
| ERROR | 81367.9 | 154 | 528.36 |  |  |
| TOTAL | 110209.9 | 161 |  |  |  |

As LOAD is strongly dependent on age, we further assessed if the odor-evoked intrinsic responses revealed an aging effect. APOE4.TREM2 mice show an increasing trend of vascular hyperresponsiveness to odor stimuli with age, while the WT mice show no significant BV changes with increasing age in the 1SD-based ROI responses (Fig. 11a, APOE4.TREM2: -5.5%_AUC_/mo, with an r^2^=0.29; WT: +2.3%_AUC_/mo, and r^2^=0.09). This is also evident in the artery-like 3SD ROI responses (Fig. 11b) and the dOB-wide rectangular ROI responses (Fig. 11c). These data emphasize not only the vascular dysfunction seen in the APOE4.TREM2 mice, but further demonstrate that the dysfunction is age dependent. These data validate this model’s translatability in studying the progression of LOAD.

**Figure 11.**
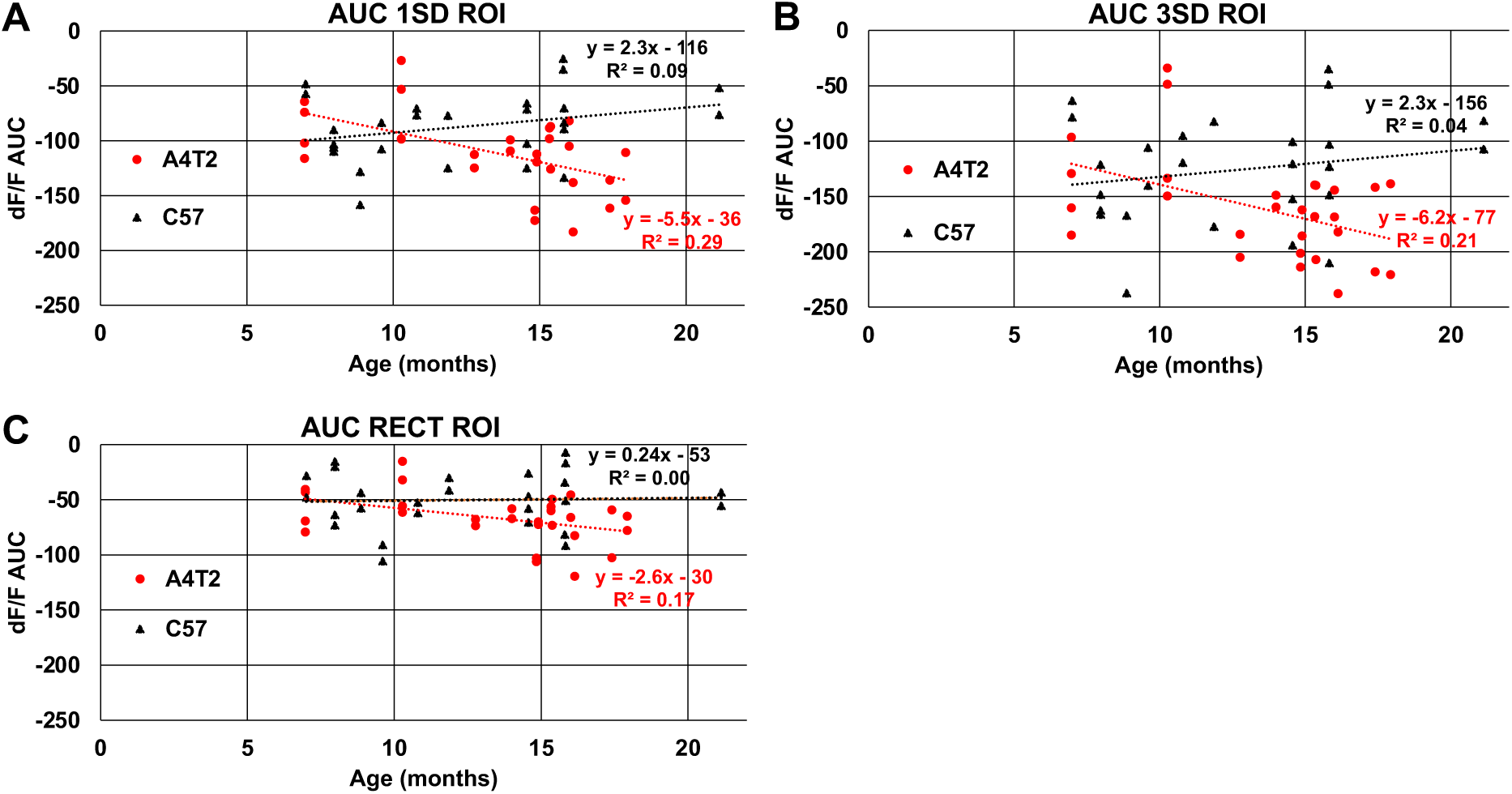
Blood volume responses in the Olfactory Bulb increase with age only in APOE4.TREM2 model mice. Scatter plots of individual mice representing intrinsic responses (AUC) for averaged 3% EB and averaged 3% MV of WT (black) and APOE4.TREM2 (red) mice by age. (**A**) Comparison for 1SD threshold ROI. (**B**) Comparison for 3SD threshold ROI. (**C**) Comparison for rectangular dOB-wide ROI.

## DISCUSSION

Resolution for the lack of efficient early diagnostic methods for LOAD first requires the validation of an effective animal model. Apoe4 and Trem2 p.R47H have been identified as two powerful genetic risk factors in the human population for LOAD, making these two strong candidates in the development of preclinical animal models. Our results suggest that the APOE4.TREM2 LOAD mouse line by the MODEL-AD consortium shows age-dependent behavioral deficits in hydration (lower fluid intake), odor-based navigation-foraging (cookie test), anxiety (open field test), and spatial memory (Morris Water Maze). In addition, we report age-dependent increases in odor induced hemodynamic responses compared to WT.

One study compared APOE4 to APOE3 in brain cells from sAD induced pluripotent stem cells (iPSCs) and revealed AD-related pathologies, such as APOE4 neurons exhibiting elevated Aβ_42_ secretions, APOE4 astrocytes having impaired Aβ uptake, and APOE4 microglia-like cells with reduced Aβ phagocytosis, relative to isogenic APOE3 cells [10].

Trem2 p.R47H has been linked to increased brain concentrations of Tumor Necrosis Factor alpha (TNF-α), heightened glutamatergic transmission, and suppressed Long-Term Potentiation (LTP) in rats. Neutralizing TNF-α with anti-TNF-α antibodies mitigated these effects on glutamatergic transmission and LTP, highlighting a role for Trem2 p.R47H in LOAD neuronal dysfunction. Interestingly, Aβ levels remained within normal range despite these alterations, suggesting evidence of early, detectable AD pathogenesis independent of the traditional Aβ pathway associated with later stages of disease [42].

While the APOE4.TREM2 mice show clear evidence of the LOAD phenotype, this current model has limitations concerning the introduction of R47H, thereby inadvertently reducing Trem2 expression and transcription. Trem2 functional loss was modeled comparing decreased expression by siRNA and RNAi-induced knockdown to the R47H variant in mice. Each displayed similar phenotypes involving decreased expression of Trem2 in microglia, decreased density of microglia in the hippocampus, and impaired downstream functionality of Trem2 mediated anti-inflammatory signaling [43]. Nonetheless, this APOE4.TREM2 model has provided useful insights into AD pathology and has served as a foundation to developing MODEL-AD [7]. Current models have corrected the cryptic splicing, attaining more accurate Trem2 expression for biologically relevant plaque observations [44].

Our data complements prior extensive results of the MODEL-AD group [7] by finding a more robust AD phenotype, likely by having chosen a complementary LOAD-targeted set of tests. Whereas Kotredes et al. reported only small behavioral deficits and brain-associated metabolic changes [7], here we demonstrate that APOE4.TREM2 mice show both robust behavioral deficits in several tasks and exacerbated intrinsic responses to odorants in the olfactory bulb.

Our sucrose preference task showed no statistical indicators of genotype-dependent anhedonia, and no impairments were found in nestlet building or the sleep/wake cycle. Genotype-dependent increased anxiety was observed in APOE4.TREM2 mice in the open field test. Using PET and autoradiography, Kotredes et al. revealed significant modifications in glycolysis and brain perfusion within regions linked to sensory integration, cognition, vision, and motor function in both B6.APOE4 and B6.APOE4.Trem2*R47H mice, compared to control mice. Female mice exhibited altered glycolysis at 4, 8, and 12 months, while male mice showed a transient hypoglycolytic phenotype at 8 months, that was virtually mitigated by 12 months. However, brain perfusion was significantly lower in both sexes by 12 months [7]. This aligns with prior studies claiming greater risk in females expressing APOE4 and diminished microglial plaque regulation [45]. Our intrinsic analysis identified robust evidence of age and genotype dependent vascular hyperresponsiveness in the dOB. All odor concentrations and ROIs revealed vascular hyperresponsiveness to both odor stimuli as a direct effect of genotype and age. Although we did intend to explore sex effects in both studies, our breeding program unfortunately did not yield sufficiently high numbers of each sex for robust exploration of sex effects, especially after taking experimentally unsuccessful trials into account (**Table 1-3 and 5**).

The APOE4.TREM2 mouse can be considered a useful model for human LOAD, developing several LOAD-related early neuro-behavioral deficits associated with the two most prevalent LOAD genetic risks. Functional MRI (fMRI) has become a valuable tool in studying how APOE-ε4 may influence brain function before clinical symptoms of AD manifest. fMRI allows researchers to examine blood flow changes in the brain that occur due to neural activity, which can provide insights into the functional connectivity (FC) of various brain regions in individuals who are genetically predisposed to AD. With the intrinsic contrast of blood oxygenation in fMRI, spatiotemporal characteristics of brain oxygenation changes are mapped in non-invasively. Both task-based (T-fMRI) and resting-state fMRI (R-fMRI) studies identified physiological changes with healthy human aging [46, 47]. FC measurements using R-fMRI technique is employed in early AD detection showing FC reductions in specific brain areas (i.e. the default mode network or DMN) [48–51]. Disruptions in DMN connectivity measured by R-fMRI indicate early AD confirmed by amyloid plaque accumulation in asymptomatic or minimally symptomatic subjects [52, 53]. Although amyloid plaque accumulation is among the early events of AD pathology, it is not directly associated with cognitive impairment [54–58]. The complexity of the relationship between the apoE genotype and FC changes highlighted by Verghese et al. [59] underscores the challenges in dissecting the mechanisms that underlie these associations. This uncertainty primarily revolves around whether the observed FC changes in APOE4 carriers are directly attributable to neural network alterations or if they are secondary effects resulting from other pathological processes, such as impaired neurovascular coupling.

Impairments of the FC in the olfactory network have been shown in early-stage AD patients and are also present in patients with MCI [50, 60]. Odor-induced fMRI activation in normal aging and AD subjects [61–63] concluded that olfactory detection tasks may serve as an early LOAD discovery tool with potential for early interventions in LOAD risk subjects [62, 64, 65].

fMRI-BOLD has been instrumental in advancing our understanding of brain activity in real time, particularly in the context of LOAD. One intriguing observation in LOAD research is the phenomenon of BOLD hyperresponsiveness, where certain brain regions exhibit increased activation compared to normal aging controls during cognitive tasks [66]. This hyperactivation is particularly notable during the early stages of AD and in individuals at high risk for developing LOAD [67]. Increased fMRI response in specific brain areas might serve as an early biomarker for LOAD, particularly in asymptomatic individuals or those who carry genetic risk factors like APOE4. Dysregulation in neurovascular coupling the relationship between local neural activity and subsequent blood flow changes might explain the observed hyperresponsiveness. In the early stages of AD there might still be sufficient vascular response to support increased metabolic demands during heightened neural activity [68].

Our robust evidence of age and genotype dependent vascular hyperresponsiveness, together with other studies reporting fMRI BOLD hyperresponsiveness during LOAD development, likely as compensatory mechanism to increasing brain dysfunction, hold promise for the use of fMRI BOLD response imaging as a tool for early diagnosis of LOAD. One study reported significant correlation in subjects’ BOLD signals within the primary olfactory cortex (POC) with their University of Pennsylvania Smell Identification Test (UPSIT), Mini-Mental State Examination (MMSE), the Mattis Dementia Rating Scale-2 (DRS-2), and Clinical Dementia Rating Scale (CDR) scores at the lowest odorant concentrations. Here BOLD signal intensity and activation volume increased significantly as a function of odorant concentration only in the AD group [69]. Overall, the current results validate the usefulness of the APOE4.TREM2 mouse model to preclinical LOAD, as well as the potential for olfactory-stimulated fMRI as an effective method in preclinical LOAD diagnostics.

## References

1. 1. Kumar, A., et al., Alzheimer Disease, in StatPearls. 2023, StatPearls Publishing Copyright © 2023, StatPearls Publishing LLC.: Treasure Island (FL).

2. Cacace, R., K. Sleegers, and C. Van Broeckhoven, Molecular genetics of early-onset Alzheimer’s disease revisited. Alzheimers Dement, 2016. 12(6): p. 733–48.

3. Vitek, M.P., et al., Translational animal models for Alzheimer’s disease: An Alzheimer’s Association Business Consortium Think Tank. Alzheimers Dement (N Y), 2020. 6(1): p. e12114.

4. Oblak, A.L., et al., Model organism development and evaluation for late-onset Alzheimer’s disease: MODEL-AD. Alzheimers Dement (N Y), 2020. 6(1): p. e12110.

5. Bird, T.D., Genetic aspects of Alzheimer disease. Genet Med, 2008. 10(4): p. 231–9.

6. Verheijen, J. and K. Sleegers, Understanding Alzheimer Disease at the Interface between Genetics and Transcriptomics. Trends Genet, 2018. 34(6): p. 434–447.

7. Kotredes, K.P., et al., Uncovering Disease Mechanisms in a Novel Mouse Model Expressing Humanized APOEepsilon4 and Trem2*R47H. Front Aging Neurosci, 2021. 13: p. 735524.

8. Lin, A.L., et al., APOE genotype-dependent pharmacogenetic responses to rapamycin for preventing Alzheimer’s disease. Neurobiol Dis, 2020. 139: p. 104834.

9. Corder, E.H., et al., Gene dose of apolipoprotein E type 4 allele and the risk of Alzheimer’s disease in late onset families. Science, 1993. 261(5123): p. 921–3.

10. Tai, L.M., et al., APOE-modulated Abeta-induced neuroinflammation in Alzheimer’s disease: current landscape, novel data, and future perspective. J Neurochem, 2015. 133(4): p. 465–88.

11. Biundo, F., et al., Interaction of ApoE3 and ApoE4 isoforms with an ITM2b/BRI2 mutation linked to the Alzheimer disease-like Danish dementia: Effects on learning and memory. Neurobiol Learn Mem, 2015. 126: p. 18–30.

12. Zheng, H., et al., Opposing roles of the triggering receptor expressed on myeloid cells 2 and triggering receptor expressed on myeloid cells-like transcript 2 in microglia activation. Neurobiol Aging, 2016. 42: p. 132–41.

13. Sudom, A., et al., Molecular basis for the loss-of-function effects of the Alzheimer’s disease-associated R47H variant of the immune receptor TREM2. J Biol Chem, 2018. 293(32): p. 12634–12646.

14. Cheng-Hathaway, P.J., et al., The Trem2 R47H variant confers loss-of-function-like phenotypes in Alzheimer’s disease. Mol Neurodegener, 2018. 13(1): p. 29.

15. Song, W., et al., Alzheimer’s disease-associated TREM2 variants exhibit either decreased or increased ligand-dependent activation. Alzheimers Dement, 2017. 13(4): p. 381–387.

16. Lin, A.L., D.A. Butterfield, and A. Richardson, mTOR: Alzheimer’s disease prevention for APOE4 carriers. Oncotarget, 2016. 7(29): p. 44873–44874.

17. Lee, J., et al., Neuroimaging Biomarkers of mTOR Inhibition on Vascular and Metabolic Functions in Aging Brain and Alzheimer’s Disease. Front Aging Neurosci, 2018. 10: p. 225.

18. Pause, B.M., et al., Perspectives on episodic-like and episodic memory. Front Behav Neurosci, 2013. 7: p. 33.

19. Mega, M.S., et al., The spectrum of behavioral changes in Alzheimer’s disease. Neurology, 1996. 46(1): p. 130–5.

20. Musiek, E.S., D.D. Xiong, and D.M. Holtzman, Sleep, circadian rhythms, and the pathogenesis of Alzheimer disease. Exp Mol Med, 2015. 47: p. e148.

21. Devanand, D.P., et al., Depressed mood and the incidence of Alzheimer’s disease in the elderly living in the community. Arch Gen Psychiatry, 1996. 53(2): p. 175–82.

22. Strauss, M.E. and P.K. Ogrocki, Confirmation of an association between family history of affective disorder and the depressive syndrome in Alzheimer’s disease. Am J Psychiatry, 1996. 153(10): p. 1340–2.

23. Doty, R.L., Olfactory dysfunction in Parkinson disease. Nat Rev Neurol, 2012. 8(6): p. 329–39.

24. Yoon, J.A., et al., Neural Compensatory Response During Complex Cognitive Function Tasks in Mild Cognitive Impairment: A Near-Infrared Spectroscopy Study. Neural Plasticity, 2019. 2019: p. 7845104.

25. Pandey, R.S., et al., Differential splicing of neuronal genes in a Trem2*R47H mouse model mimics alterations associated with Alzheimer’s disease. BMC Genomics, 2023. 24(1): p. 172.

26. Knouff, C., et al., Apo E structure determines VLDL clearance and atherosclerosis risk in mice. J Clin Invest, 1999. 103(11): p. 1579–86.

27. Coronas-Samano, G., A.V. Ivanova, and J.V. Verhagen, The Habituation/Cross-Habituation Test Revisited: Guidance from Sniffing and Video Tracking. Neural Plast, 2016. 2016: p. 9131284.

28. Romano, A., et al., Depressive-like behavior is paired to monoaminergic alteration in a murine model of Alzheimer’s disease. Int J Neuropsychopharmacol, 2015. 18(4).

29. Van Dam, D., et al., Age-dependent cognitive decline in the APP23 model precedes amyloid deposition. Eur J Neurosci, 2003. 17(2): p. 388–96.

30. Chen, Y., et al., A non-transgenic mouse model (icv-STZ mouse) of Alzheimer’s disease: similarities to and differences from the transgenic model (3xTg-AD mouse). Mol Neurobiol, 2013. 47(2): p. 711–25.

31. Galeano, P., et al., Longitudinal analysis of the behavioral phenotype in a novel transgenic rat model of early stages of Alzheimer’s disease. Front Behav Neurosci, 2014. 8: p. 321.

32. Morris, R., Developments of a water-maze procedure for studying spatial learning in the rat. J Neurosci Methods, 1984. 11(1): p. 47–60.

33. Javed, H., et al., S-allyl cysteine attenuates oxidative stress associated cognitive impairment and neurodegeneration in mouse model of streptozotocin-induced experimental dementia of Alzheimer’s type. Brain Res, 2011. 1389: p. 133–42.

34. Deacon, R.M., Assessing nest building in mice. Nat Protoc, 2006. 1(3): p. 1117–9.

35. Wesson, D.W. and D.A. Wilson, Age and gene overexpression interact to abolish nesting behavior in Tg2576 amyloid precursor protein (APP) mice. Behav Brain Res, 2011. 216(1): p. 408–13.

36. Donohue, K.D., et al., Assessment of a non-invasive high-throughput classifier for behaviours associated with sleep and wake in mice. Biomed Eng Online, 2008. 7: p. 14.

37. Duncan, M.J., et al., Effects of aging and genotype on circadian rhythms, sleep, and clock gene expression in APPxPS1 knock-in mice, a model for Alzheimer’s disease. Exp Neurol, 2012. 236(2): p. 249–58.

38. Mang, G.M., et al., Evaluation of a piezoelectric system as an alternative to electroencephalogram/ electromyogram recordings in mouse sleep studies. Sleep, 2014. 37(8): p. 1383–92.

39. Sethi, M., et al., Increased fragmentation of sleep-wake cycles in the 5XFAD mouse model of Alzheimer’s disease. Neuroscience, 2015. 290: p. 80–9.

40. Ratzlaff, E.H. and A. Grinvald, A tandem-lens epifluorescence macroscope: hundred-fold brightness advantage for wide-field imaging. J Neurosci Methods, 1991. 36(2-3): p. 127–37.

41. Crawley, J. and F.K. Goodwin, Preliminary report of a simple animal behavior model for the anxiolytic effects of benzodiazepines. Pharmacol Biochem Behav, 1980. 13(2): p. 167–70.

42. Ren, S., et al., Microglia TREM2R47H Alzheimer-linked variant enhances excitatory transmission and reduces LTP via increased TNF-α levels. eLife, 2020. 9: p. e57513.

43. Liu, W., et al., Trem2 promotes anti-inflammatory responses in microglia and is suppressed under pro-inflammatory conditions. Hum Mol Genet, 2020. 29(19): p. 3224–3248.

44. Tran, K.M., et al., A Trem2(R47H) mouse model without cryptic splicing drives age- and disease-dependent tissue damage and synaptic loss in response to plaques. Mol Neurodegener, 2023. 18(1): p. 12.

45. Stephen, T.L., et al., APOE genotype and sex affect microglial interactions with plaques in Alzheimer’s disease mice. Acta Neuropathologica Communications, 2019. 7(1): p. 82.

46. Buckner, R.L., J.R. Andrews-Hanna, and D.L. Schacter, The brain’s default network: anatomy, function, and relevance to disease. Ann N Y Acad Sci, 2008. 1124: p. 1–38.

47. D’Esposito, M., et al., The effect of normal aging on the coupling of neural activity to the bold hemodynamic response. Neuroimage, 1999. 10(1): p. 6–14.

48. Damoiseaux, J.S., et al., Functional connectivity tracks clinical deterioration in Alzheimer’s disease. Neurobiol Aging, 2012. 33(4): p. 828 e19-30.

49. Greicius, M.D., et al., Default-mode network activity distinguishes Alzheimer’s disease from healthy aging: evidence from functional MRI. Proc Natl Acad Sci U S A, 2004. 101(13): p. 4637–42.

50. Lu, J., et al., Disruptions of the olfactory and default mode networks in Alzheimer’s disease. Brain Behav, 2019. 9(7): p. e01296.

51. Mousa, D., N. Zayed, and I.A. Yassine, Alzheimer disease stages identification based on correlation transfer function system using resting-state functional magnetic resonance imaging. PLoS One, 2022. 17(4): p. e0264710.

52. Pacyna, R.R., et al., Rapid olfactory decline during aging predicts dementia and GMV loss in AD brain regions. Alzheimers Dement, 2022.

53. Sperling, R.A., et al., Amyloid deposition is associated with impaired default network function in older persons without dementia. Neuron, 2009. 63(2): p. 178–88.

54. Aizenstein, H.J., et al., Frequent amyloid deposition without significant cognitive impairment among the elderly. Arch Neurol, 2008. 65(11): p. 1509–17.

55. Buckner, R.L., et al., Molecular, structural, and functional characterization of Alzheimer’s disease: evidence for a relationship between default activity, amyloid, and memory. J Neurosci, 2005. 25(34): p. 7709–17.

56. Mormino, E.C. and K.V. Papp, Amyloid Accumulation and Cognitive Decline in Clinically Normal Older Individuals: Implications for Aging and Early Alzheimer’s Disease. J Alzheimers Dis, 2018. 64(s1): p. S633–S646.

57. Morris, G.P., I.A. Clark, and B. Vissel, Inconsistencies and controversies surrounding the amyloid hypothesis of Alzheimer’s disease. Acta Neuropathol Commun, 2014. 2: p. 135.

58. Palmqvist, S., et al., Earliest accumulation of beta-amyloid occurs within the default-mode network and concurrently affects brain connectivity. Nat Commun, 2017. 8(1): p. 1214.

59. Verghese, P.B., J.M. Castellano, and D.M. Holtzman, Apolipoprotein E in Alzheimer’s disease and other neurological disorders. Lancet Neurol, 2011. 10(3): p. 241–52.

60. Lu, J., et al., Functional Connectivity between the Resting-State Olfactory Network and the Hippocampus in Alzheimer’s Disease. Brain Sci, 2019. 9(12).

61. Feng, Q., et al., Objective Assessment of Hyposmia in Alzheimer’s Disease From Image and Behavior by Combining Pleasant Odor With Unpleasant Odor. Front Neurol, 2021. 12: p. 697487.

62. Steffener, J., et al., Odorant-induced brain activation as a function of normal aging and Alzheimer’s disease: A preliminary study. Behav Brain Res, 2021. 402: p. 113078.

63. Zhang, H., et al., Olfactory fMRI Activation Pattern Across Different Concentrations Changes in Alzheimer’s Disease. Front Neurosci, 2019. 13: p. 786.

64. Bell, S.A., et al., Development of novel measures for Alzheimer’s disease prevention trials (NoMAD). Contemp Clin Trials, 2021. 106: p. 106425.

65. Motter, J.N., et al., Odor identification impairment and cholinesterase inhibitor treatment in Alzheimer’s disease. Alzheimers Dement (Amst), 2021. 13(1): p. e12158.

66. Lockwood, C.T. and C.J. Duffy, Hyperexcitability in Aging Is Lost in Alzheimer’s: What Is All the Excitement About? Cereb Cortex, 2020. 30(11): p. 5874–5884.

67. Targa Dias Anastacio, H., N. Matosin, and L. Ooi, Neuronal hyperexcitability in Alzheimer’s disease: what are the drivers behind this aberrant phenotype? Transl Psychiatry, 2022. 12(1): p. 257.

68. Shabir, O., J. Berwick, and S.E. Francis, Neurovascular dysfunction in vascular dementia, Alzheimer’s and atherosclerosis. BMC Neurosci, 2018. 19(1): p. 62.

69. Wang, J., et al., Olfactory deficit detected by fMRI in early Alzheimer’s disease. Brain Res, 2010. 1357: p. 184–94.

